# Simulation and quantitative analysis of spatial centromere distribution patterns

**DOI:** 10.1101/2025.01.22.634320

**Authors:** Adib Keikhosravi, Krishnendu Guin, Gianluca Pegoraro, Tom Misteli

## Abstract

A prominent feature of eukaryotic chromosomes are centromeres, which are specialized regions of repetitive DNA required for faithful chromosome segregation during cell division. In interphase cells centromeres are non-randomly positioned in the three-dimensional space of the nucleus in a cell-type specific manner. The functional relevance and the cellular mechanisms underlying this observation are unknown, and quantitative methods to measure distribution patterns of centromeres in 3D space are needed. Here we have developed an analytical framework that combines robust clustering metrics and advanced modeling techniques for the quantitative analysis of centromere distributions at the single cell level. To identify a robust quantitative measure for centromere clustering, we benchmarked six metrics for their ability to sensitively detect changes in centromere distribution patterns from high-throughput imaging data of human cells, both under normal conditions and upon experimental perturbation of centromere distribution. We find that Ripley’s K Score has the highest accuracy with minimal sensitivity to variations in centromeres number, making it the most suitable metric for measuring centromere distributions. As a complementary approach, we also developed and validated spatial models to replicate centromere distribution patterns, and we show that a radially shifted Gaussian distribution best represents the centromere patterns seen in human cells. Our approach creates tools for the quantitative characterization of spatial centromere distributions with applications in both targeted studies of centromere organization as well as in unbiased screening approaches.

## Introduction

The spatial distribution of many nuclear compartments, including proteinaceous bodies and genomic domains, is non-random (Misteli 2020). Gene-rich chromosomes, for example, tend to occupy central nuclear regions, whereas gene-poor chromosomes are typically more peripheral (Kreth et al. 2004). Similarly, transcription sites and replicating loci, often associated with active chromatin, are often more centrally localized in the cell nucleus, whereas inactive heterochromatin is generally positioned at the nuclear periphery or in the proximity of nucleoli (Hu et al. 2008; Faro-Trindade and Cook 2006; Osborne et al. 2007). The spatial organization of chromosomes within the nucleus has been suggested to be integral to cellular function, impacting processes such as chromatin accessibility, gene expression, DNA repair, and DNA replication (Misteli 2020).

Centromeres are specialized regions on each chromosome that are crucial for accurate chromosome segregation. During cell division, centromeres serve as the attachment point for the microtubule spindle via the kinetochore protein complex (McKinley and Cheeseman 2016; Fukagawa and Earnshaw 2014). Like other nuclear structures, centromeres assume an apparently non-random spatial distribution within the nucleus (Sullivan and Sullivan 2020; Sullivan et al. 2011; Solovei et al. 2004; M. Cremer et al. 2001), and in a cell-type specific manner (Solovei et al. 2004; Alcobia, Dilão, and Parreira 2000). For example, in human stem cells, the majority of centromeres clusters strongly near the nucleoli (Bersaglieri et al. 2022; Rodrigues et al. 2023), and this association is weakened as the stem cells differentiate (Rodrigues et al. 2023). Similarly, the association of centromeres with the nucleolus has also been observed in other non-stem cell types (Kumar et al. 2024). Although the proximity of rDNA loci to the centromeres on all five rDNA-containing chromosome may partly account for this behavior (Altemose et al. 2022), the spatial proximity of centromeres to the nucleolus may also have functional consequences as suggested by the finding that transcription from alpha-satellite repeats of centromeres is limited near nucleoli (Bury et al. 2020), and by the observation that peripheral chromosomes have a higher rate of mis-segregation compared to the centrally positioned ones (Klaasen et al. 2022). Furthermore, variations in centromeres clustering have also been implicated in diverse biological phenomena, including development, cancer progression, and response to cellular stress (Krämer, Maier, and Bartek 2011; McKinley and Cheeseman 2016).

Little is known about the cellular factors and mechanisms that determine spatial centromere distribution. Candidate-based studies in cancer cell lines have demonstrated that alterations in centromere clustering can result from changes in chromatin organization, impacting genomic stability and mitotic fidelity (Naughton et al. 2022). Similarly, clustering variations have been associated with perturbations of the NCAPH2 subunit of the condensin II complex (Hoencamp et al. 2021), which regulates chromatin compaction and spatial genome organization (Stephens et al. 2013). However, the specific molecular mechanisms and processes that determine spatial organization of centromeres remain unknown.

High-Throughput Imaging (HTI) assays are a powerful approach to systematically study cellular phenotypes, including centromere distribution, at the single cell level, and in an unbiased fashion when paired with functional genomics perturbations, such as RNAi or CRISPR-Cas9 gene knock outs (Pegoraro and Misteli 2017). These assays are ideally suited to perform functional genomics screens to identify regulators of centromeric localization, and they rely on robust metrics to quantify centromeres clustering patterns across millions of cells and across thousands of biological conditions. Unfortunately, such metrics are currently missing.

To fill this gap, we conducted extended testing to identify novel analytical tools to quantitatively analyze centromere localization patterns in human cells, including several global and local clustering metrics and spatial distribution modeling approaches. First, we benchmarked six clustering metrics and tested multiple spot generation models to evaluate their utility in measuring different clustering patterns on simulated data. Then, we tested these metrics on HTI experimental data from multiple human cell types, and we identified a derivative version of the Ripley’s K Score as the most robust indicator of different clustering patterns in single cells. Finally, to extend the applicability of our framework, we also surveyed several modeling approaches to fit the imaging data and re-create realistic centromere distributions *in silico*. We tested these models on multiple human cell lines, demonstrating their versatility and accuracy in capturing diverse centromere localization patterns and after experimental perturbation of centromere clustering patterns. Our approach establishes quantitative tools for the study of centromere localization and function they have potential for wider application in genome research.

## Methods

### siRNA Oligos Transfection and Immunofluorescence

The image dataset containing human colon cancer HCT116-Cas9 cells reverse-transfected with siRNA oligos against the *NCAPH2* gene and scrambled negative siRNA control in 384-well plates has been described (Keikhosravi et al. 2024).

### Cell Growth and Centromere Visualization

All cells were grown in 384-well plates (CellVis, Cat. No. P384-1.5H-N). The sources of cell lines, culture conditions, media compositions and relevant references of respective culture protocols are described in Supplementary Table 1. Cell lines were grown for 72 hours after cell seeding before fixation, except for the iPS WTC11 cells that were grown for 3-5 days and fixed before the colony edges merged with each other. All cell lines were fixed with 2% paraformaldehyde (PFA, Electron Microscopy Sciences, Cat. No. 15710) solution in media by adding one equal volume of 4% PFA solution in PBS to the cell growth medium in each well for 15 minutes at room temperature. Fixed cells were then subjected to immunofluorescent (IF) staining using anti-CENP-C antibody (#PD030, MBL Co. Ltd.) and DAPI (4′,6-diamidino-2-phenylindole) (Thermo Fisher Scientific, Cat. No. 62248) as previously described (Keikhosravi et al. 2024). The total number of cells analyzed from this dataset are: A375, n=11,925; MDA-MB-231, n=6,347; HFF-hTERT, n=1,322; hTERT-RPE1, n=3,641; A549, n=7,786; WTC11, n=6,071; HAP1, n=18,359; and HCT116, n=16,116.

### High-Throughput Image Acquisition

For IF experiments, images were acquired using 405 nm (DAPI channel) or 488 nm (CENP-C channel) excitation lasers and a 405/488/561/640 nm excitation dichroic mirror. A 60X water-immersion objective (NA 1.2) was employed, paired with 445/45 nm (DAPI channel) or 525/50 nm (CENP-C channel) bandpass emission filters. A 16-bit sCMOS camera (2048 × 2048 pixels, 1×1 binning, pixel size: 0.108 microns) was used for the capture of image Z-stacks spanning 14 microns in depth, collected at 1-micron intervals and maximally projected in on the fly. Images were acquired from 22 fields of view (FOV) per well.

### High-Throughput Image Analysis

The analysis of imaging data was carried out using HiTIPS, a high-throughput image analysis software designed to analyze cell-based assays in fixed and live cells as previously described (Keikhosravi et al. 2024). Maximally projected DAPI images were used for nucleus segmentation, while CENP-C images were used for CENP-C spot finding and localization (Figure 1). Specific analysis parameters were selected in HiTIPS tailored to align with the average nucleus size, as well as the size and brightness of the centromere spots observed. The GPU-based CellPose algorithm for nuclei segmentation (Pachitariu and Stringer 2022) was used in conjunction with the Laplacian of Gaussian method for spot detection. Spot positions were determined as the center of gravity of the segmented spot.

**Figure 1.**
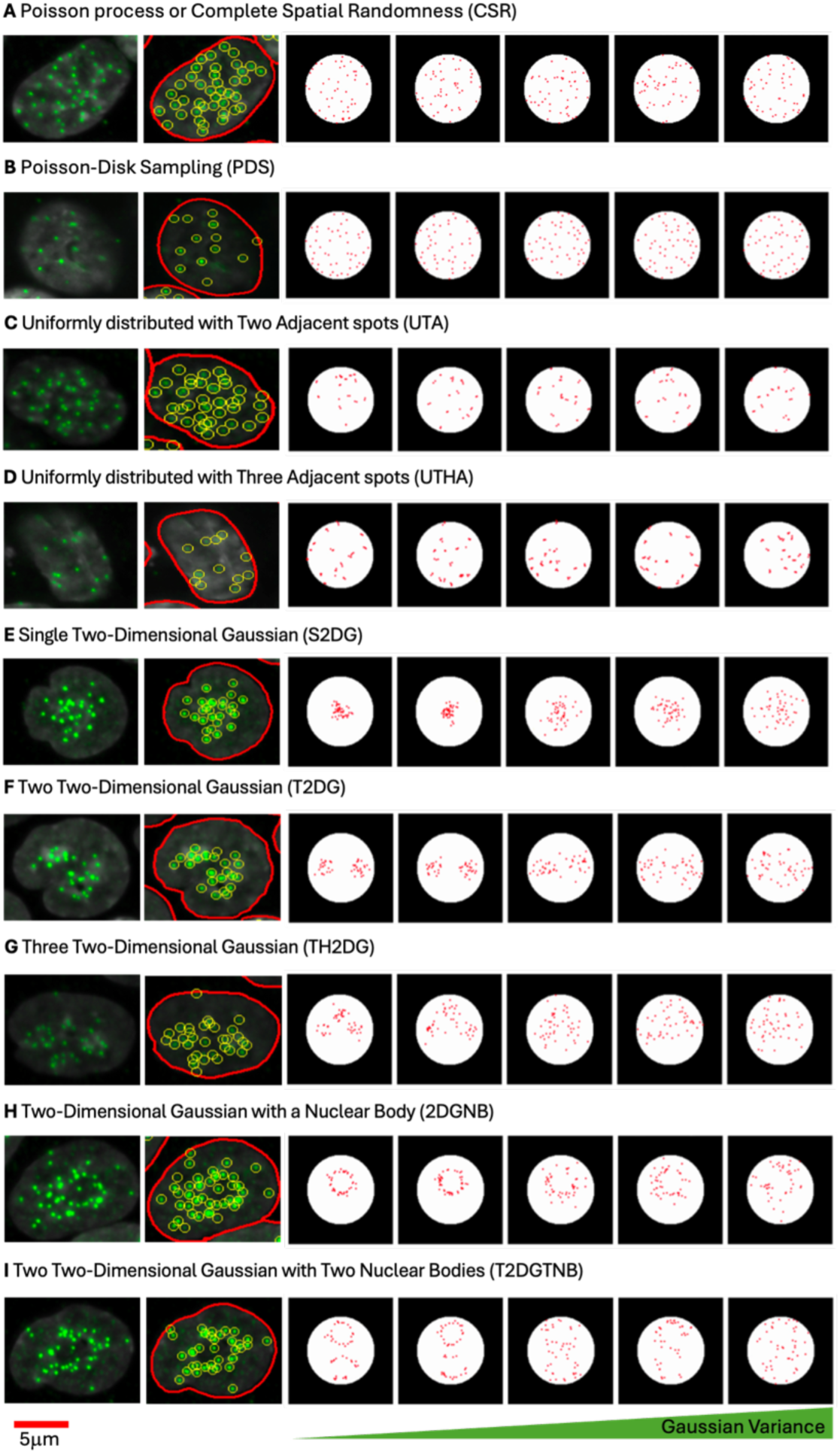
Simulated spatial distribution patterns of centromeres modeled to reflect various clustering and dispersion behaviors observed in HCT116-Cas9 colorectal cancer cells. For each row, the two leftmost images display the fluorescence image of centromere protein CENP-C in a HCT116 cell (green), followed by extracted spot locations (yellow circles) outlined within the boundary to indicate the nucleus outline (red). Each row represents a distinct spatial distribution model: (A) Complete Spatial Randomness (CSR), where centromeres are uniformly and randomly distributed; (B) Poisson-Disk Sampling (PDS), enforcing a minimum distance between spots to prevent overlap; (C) and (D) Uniformly distributed centromeres with two (UTA) or three (UTHA) adjacent spots mimicking small clusters; (E–G) Single (S2DG), Two (T2DG), or Three (TH2DG) Two-Dimensional Gaussian distributions, representing varying levels of clustering within larger nuclear regions; (H) Two-Dimensional Gaussian with a Nuclear Body (2DGNB), excluding the nuclear body area; and (I) Two Two-Dimensional Gaussian distributions with Two Nuclear Bodies (T2DGTNB). For simulated distributions, a gradient of Gaussian Variance increases from left to right as indicated.

### Methods for generating synthetic spot patterns

To evaluate centromere spot pattern characterization metrics, synthetic images containing simulated distributions of spots were generated under controlled spatial arrangements within a circular region representing the cell nucleus. The synthetic spots were generated on an image patch of pixels. A circular region was defined at the center of this patch, with a radius pixels and center coordinates. All synthetic spots were constrained within this circular region, ensuring consistency across pattern types. The constraint for any spot was: (*x* − *c_x_*)^2^ + (*y* − *c_y_*)^2^ ≤ *r*^2^. Each distribution was designed with specific statistical properties to mimic potential centromere clustering patterns.

#### Poisson Process or Complete Spatial Randomness (CSR)

To generate a uniform distribution of spots, random points were sampled according to a uniform distribution within the bounds [*c_x_* − *r*, *c_x_* + *r*] for *x* and [*c_y_* − *r*, *c_y_* + *r*] for *y*. Each point was retained only if it laid within the circular region as defined above. For each sample, 46 spots were generated. This uniform sampling, which is also referred to as Poisson process, provided a baseline distribution to assess other pattern types against a spatially random background.

#### Single Two-Dimensional Gaussian (S2DG)

For clustering patterns, spots were generated based on a 2D Gaussian distribution centered at (128,128) with varying covariance matrices. The covariance matrices Σ were systematically varied in size (from 50 to 1000 pixels squared in increments) and orientation (from 0 to *π* radians).

The covariance matrix for each Gaussian distribution was defined as:

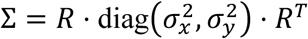

where *R* is the rotation matrix for angle *θ*, and *σ_x_*and *σ_y_*represent the standard deviations along the rotated principal axes. For each sample, points were drawn from the Gaussian distribution, and only the first 46 spots within the circular boundary were retained. This process provided various levels of spot clustering within the circle.

#### Two (T2DG) and Three (TH2DG) Two-Dimensional Gaussian

To models for multi-modal clustering, Gaussian Mixture Models (GMMs) (Reynolds 2009), were used with either two or three Gaussian components (modes) centered within the circular region. For the two-mode GMM, the component centers were positioned symmetrically around (*c_x_*, *c_y_*) at distances of 25 pixels, representing two distinct clusters. For the three-mode GMM, three Gaussian components were similarly spaced 20 pixels apart in a triangular arrangement around (*c_x_*, *c_y_*).

Each component’s covariance matrix was systematically varied using the same range of sizes and orientations as described for the single Gaussian. Samples from the GMM were filtered to retain only those within the circle. The GMM model used equal weights for each component:

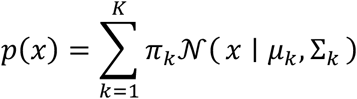

where *K* is the number of modes (2 or 3), and *π_k_* is the component weight (0.5 for 2-mode, 1/3 for 3-mode).

- Means:

- 2-Mode: (128,103) and (128,153)
- 3-Mode: (128,103), (128,153), and (103,128)
- Covariance range: Size from 50 to 1000, orientations from 0 to *π*

For each sample, points were drawn from the corresponding distribution, and only the first 46 spots within the circular boundary were retained.

#### Poisson-Disk Sampling (PDS)

A Poisson-disk sampling approach (Wang 2024) was applied to generate dispersed spots, enforcing a minimum inter-spot distance of 10 pixels. This approach simulated highly dispersed patterns, characteristic of spatially non-clustered arrangements. Candidate points were generated uniformly within the circular boundary, but each new point was retained only if it satisfied the minimum distance requirement from all existing spots. This technique was applied to generate 46 spots per sample while maintaining a minimum spacing constraint.

#### Uniformly distributed with Two (UTA) and Three (UTHA) Adjacent spots

To model proximity effects, two and three adjacent spots were generated by perturbing a subset of initial points with a random shift *ϵ* sampled from [−3,3] pixels. For two adjacent spots, the perturbation was applied to 15 spots out of the 46, generating a second, closely placed spot for each. For three adjacent spots, two perturbations were applied to 15 initial spots, creating two adjacent spots per each of the 15. The first 46 points within the circular boundary were retained after applying the perturbation.

### Spot Clustering Metrics

To analyze centromere spot patterns within synthetic distributions, we employed different metrics that capture clustering, modularity, spatial autocorrelation, and dispersion characteristics. Each metric was computed based on specific parameter settings and methodologies, as outlined below.

#### Ripley’s K Score

The Ripley’s K function 8*K*(*r*): (Ripley 1976) was used to quantify spatial clustering by calculating the difference between the expected number of neighboring points within a radius *r* of each point, and the expected number of neighboring points within a radius *r* of each point for a homogeneous spatial Poisson process, or Complete State of Randomness (CSR). The Ripley’s K Score is maximized when the expected number of spots from any given point is larger than the one for the CSR distribution for a large number of radiuses *r*. This metric results in higher values for spot patterns from one large cluster such as single 2D Gaussian with and without nuclear bodies. However, for dispersed small clusters, and even two and three mode 2D Gaussians, this Ripley’s K Score is expected to return significantly lower values. We report the clustering percentage as the fraction of radii where *K*(*r*) exceeds the Poisson expectation, indicating clustering. The calculation follows:

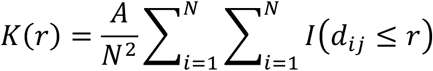

where *A* is the area of the region, *N* is the number of points, *d_ij_* is the distance between points *i* and *j*, and *I* is an indicator function that is 1 if *d_ij_* ≤ *r* and 0 otherwise. We calculate Ripley’s K clustering score, or clustering percentage, as:

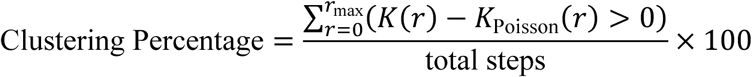

where *K*_Poisson_(*r*) is the expectation for a spatially random distribution.

#### Assortativity Coefficient

The Assortativity coefficient is a measure of connectedness of nodes with similar degrees in a graph (Shizuka and Farine 2016), and it is computed for a *k*--nearest neighbor (k-NN) graph created from the centromere points, reflecting the tendency of nodes to connect to others with similar degree (K=10). Assortativity is based on degree measures the correlation between nodes’ degrees and the degrees of their neighbors, and is defined as:

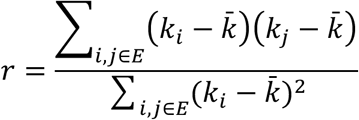

where *k_i_* and *k*_j_ are the degrees of nodes *i* and *j* in the graph, *k̄* is the mean node degree, and *E* represents the set of edges in the graph.

Assortativity is reported as a single coefficient between −1 (disassortative) and +1 (assortative). Since the node degrees are calculated as the inverse of squared distance of spots from each other, this metric is expected to be maximum when most of the spots are surrounding the nuclear bodies such as nucleolus. However, the values are still very close to zero and very closely followed by spots generated using a single 2D Gaussian distribution.

#### Modularity

Modularity *Q* measures the density of connections within clusters compared to connections between clusters (Brandes et al. 2008), reflecting how well the graph divides into modules. The Louvain algorithm was used to compute modularity, where:

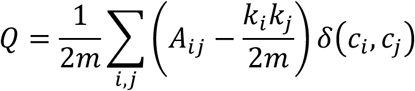

Here, *A_ij_* is the adjacency matrix, *k_i_* and *k_j_* are the degrees of nodes *i* and *j*, *m* is the total number of edges, and *δ*(*c_i_*, *c*_j_) is 1 if nodes *i* and *j* belong to the same community and 0 otherwise. The modularity index *Q* ranges from −1 to 1, with values closer to 1 indicating stronger modularity. This is expected to be maximum in presence of small clusters dispersed within nuclei.

#### Moran’s I

Moran’s I statistic was used to evaluate spatial autocorrelation of centromere spots (Tiefelsdorf and Boots 1995). Moran’s I measures whether spots are clustered (positive value), dispersed (negative value), or randomly distributed (near zero). Moran’s I is defined as:

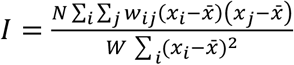

where *N* is the number of points, *x_i_* and *x*_j_ are the coordinates of points *i* and *j*, *x̅* is the mean coordinate, *w_ij_* is the spatial weight (set to 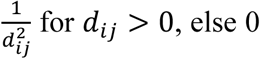), and *W* = ∑*_i_* ∑_j_ *w_ij_* is the sum of all weights. Moran’s I ranges from −1 (complete dispersion) to 1 (perfect clustering), with zero indicating no autocorrelation. Moran’s I is expected to be maximized by the presence of local clusters further from the centroid of the spots, such as uniformly distributed with two and three adjacent spots and minimized when all the spots are clustered at the center of the nucleus as generated by single 2D Gaussian distribution.

#### Mean Nearest Neighbor Distance (MNND)

Mean Nearest Neighbor Distance (MNND) quantifies the average distance to the nearest neighboring spot (Clark and Evans 1954), providing a measure of into the clustering density of the centromeres. It is calculated as:

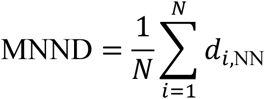

where *d_i,NN_* is the distance between point iii and its nearest neighbor and *N* is the total number of points. A lower MNND indicates tighter clustering, while a higher value suggests more dispersed points. MNND is expected to be maximized with a hard-core process spot pattern and minimized when each spot has at least one other spot in proximity.

#### Dispersion Index

The Dispersion Index quantifies the level of spatial dispersion or clustering by comparing the variance of pairwise distances to their mean distance (Myers 1978). The dispersion index *D* is defined as:

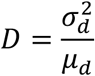

where 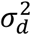 is the variance of the pairwise distances between points and *μ_d_*is the mean of those distances. A higher dispersion index indicates greater variability in distances, suggesting spatial clustering, whereas a lower index indicates uniform dispersion. The Dispersion Index is used to assess the spread of centromeres within the defined circular area, and it will be maximized when the average of pairwise distances of the spots are minimum and the variance is high. Both these conditions can occur simultaneously with uniformly distributed with two and three adjacent spots.

### Centromere Spot localization Modeling methods Using Gaussian Distributions

To model the localization of the centromere spots we used several approaches based on spot location and distribution of their pairwise distances. Below each approach is explained in detail:

#### Uniform Distribution of Spots on Cell Nucleus as Benchmarking Method

To establish a baseline for centromere spot localization, a uniform distribution of spots was generated within the boundaries of the cell nucleus that is also known as Poisson Process. This method simulates a completely random spatial arrangement, providing a benchmark for comparing clustering metrics and other spatial pattern models.

#### Modeling of Centromere Localization Using Cell Shape

To model centromere localization based on nuclear geometry, an ellipse was fitted to the segmented nucleus. The parameters of the ellipse, specifically the major axis *a* and minor axis *b*, were used to define the spatial boundaries for generating a two-dimensional Gaussian distribution. The Gaussian distribution was centered at the nucleus center (*x_c_*, *y*_*c*)_, and the probability density function at any point (*x*, *y*) was given by:

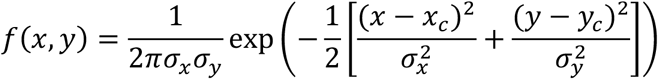

where 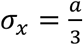 and 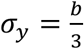 represent the standard deviations along the major and minor axes, respectively, to ensure realistic clustering within the nuclear boundary. The generated spots follow this elliptical Gaussian distribution, accurately reflecting the centromere patterns under the spatial constraints imposed by the nuclear shape.

#### Radial Distribution and Ripley’s *K*(*r*) Function Calculation

For this approach, we calculated Ripley’s *K*(*r*) function, which describes the expected number of points within a distance *r* from a randomly chosen point, normalized by the intensity *λ* of the point process. The *K*(*r*) function for our radially symmetric Gaussian distribution is defined as:

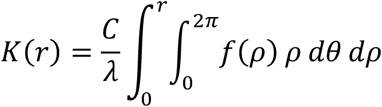

where *f*(*ρ*) is the radial probability density function (PDF) of a 2D Gaussian distribution centered at the origin, given by:

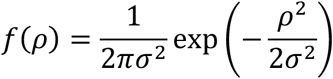

Converting to polar coordinates allows us to express the expected number of points within distance *r* by integrating *f*(*ρ*) over the radial distance *ρ* from the origin. The inner integral over *θ* simplifies due to the independence of *f*(*ρ*) from the angular coordinate, reducing *K*(*r*) to:

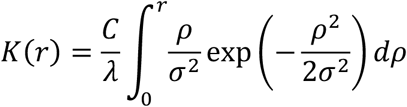

To further evaluate this expression, we used the substitution 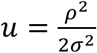, transforming the integral into:

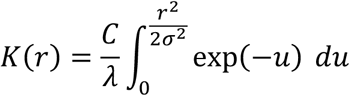

which yields:

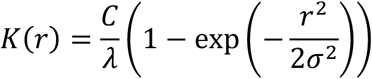

This closed-form solution provides a cumulative description of the expected clustering of points up to radius *r*, based on a Gaussian spatial distribution of centromere locations.

#### Bayesian Estimation of Radially shifted Gaussian distribution using spots coordinates

Given the observed *r*_1_, *r*_2_, …, *r_n_*, and knowing that each *r_i_* is drawn from a Gaussian distribution with mean *r*_0_ and variance *σ*^2^, the likelihood function *L*(*r*_0_, *σ*) can be expressed as:

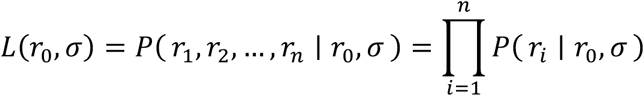

Where each *P*(*r_i_* ∣ *r*_0_, *σ*) is given by:

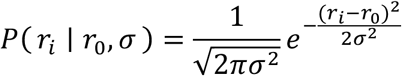

Thus, the full likelihood function becomes:

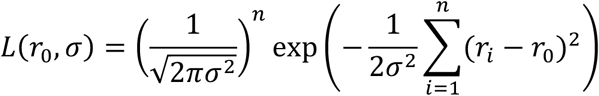

Given that *r*_0_ and *σ* are themselves normally distributed, we can define the priors:

- Prior on 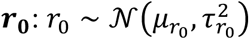, where 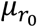 and 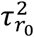 are the mean and variance of the prior on *r*_0_.

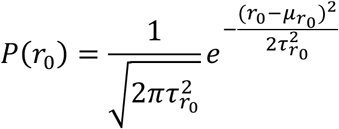

- Prior on 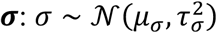, where *μ*_σ_ and 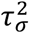 are the mean and variance of the prior on *σ*.

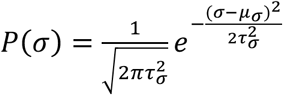

The goal is to find the posterior distribution *P*(*r*_0_, *σ* ∣ *r*_1_, *r*_2_, …, *r_n_*), which is proportional to the product of the likelihood and the prior:

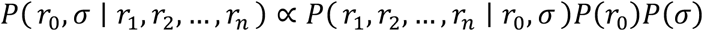

Substituting the likelihood and priors:

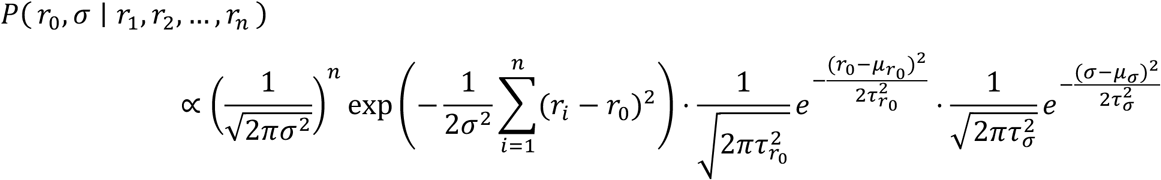

This can be expressed more compactly as:

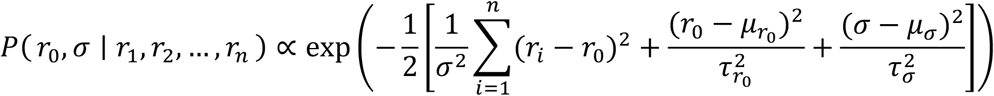

#### Bayesian Estimation of the Radially Shifted Gaussian Distribution Using Pairwise Distances

In polar coordinates, a radially shifted Gaussian distribution can be represented as:

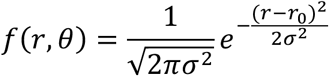

where:

- *r* is the radial distance from the origin.
- *θ* is the angular coordinate.
- *r*_0_ is the mean radius of the doughnut shape.
- σ is the standard deviation controlling the thickness of the doughnut.

We then convert the polar coordinates (*r*_1_, *θ*_1_) and (*r*_2_, *θ*_2_) into Cartesian coordinates:

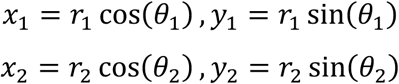

The Euclidean distance *d* between the two points (*x*_1_, *y*_1_) and (*x*_2_, *y*_2_) is given by:

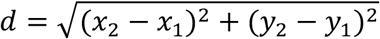

Substituting the expressions for *x*_1_, *y*_1_ and *x*_2_, *y*_2_ gives:

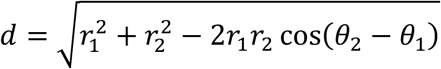

To derive the distribution of pairwise distances, we need to consider the probability distributions of *r*_1_, *r*_2_, and *θ*_2_ − *θ*_1_. Assuming *r*_1_ and *r*_2_ are independently drawn from the radially shifted Gaussian distribution and *θ*_2_ − *θ*_1_ is uniformly distributed over [0,2*π*], the distribution of *d* can be obtained by integrating over all possible values of *r*_1_, *r*_2_, and *θ*_2_ − *θ*_1_:

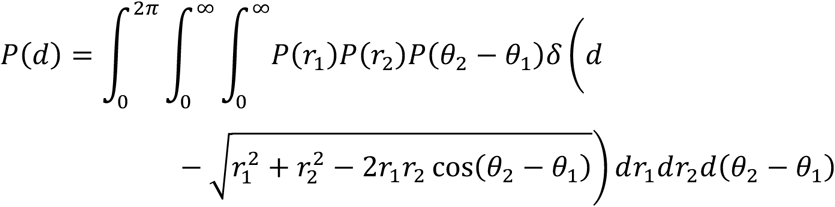

Here, *P*(*r*_1_) and *P*(*r*_2_) are the radial distributions (Gaussian with mean *r*_0_ and variance *σ*^2^), *δ* is the Dirac delta function that ensures the integration considers only valid pairwise distances. where *P*(*r*_1_) and *P*(*r*_2_) are Gaussian distributions:

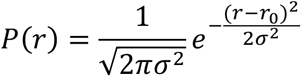

and 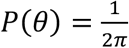 is the uniform distribution of the angle difference. Simplifying the angle integral by noting that *P*(*θ*) is uniform:

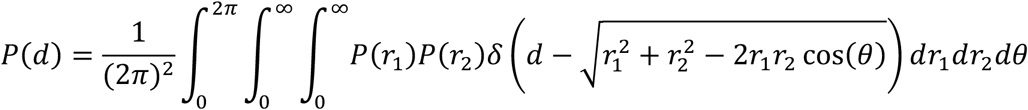

By substituting the variable *u* = cos(*θ*), we have *du* = − sin(*θ*) *dθ*. The limits of integration for *u* are from −1 to 1.

The delta function imposes the condition that: 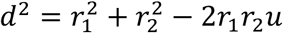. This, in turn implies: 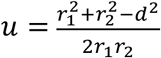. So, the integral over *u* becomes:

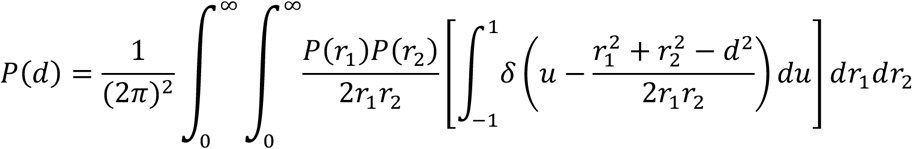

The delta function reduces the integral over *u*, giving:

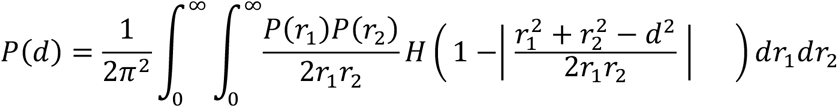

where *H*(*x*) is the Heaviside step function ensuring that 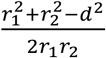 lies within [−1,1].

The final form of *P*(*d*) involves evaluating the integral:

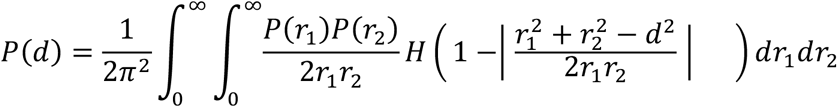

Given the observed *r*_1_, *r*_2_, …, *r_n_*, and knowing that each *r_i_* is drawn from a Gaussian distribution with mean *r*_0_ and variance *σ*^2^, the likelihood function *L*(*r*_0_, *σ*) can be expressed as:

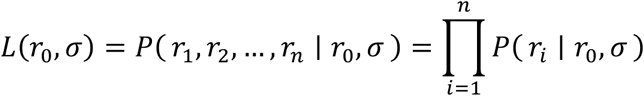

Where each *P*(*r_i_* ∣ *r*_0_, *σ*) is given by:

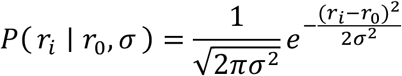

Thus, the full likelihood function becomes:

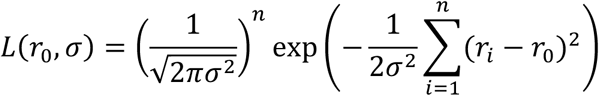

Given that *r*_0_ and *σ* are themselves normally distributed, we can define the priors:

- Prior on 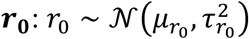, where 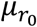 and 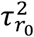 are the mean and variance of the prior on *r*_0_.

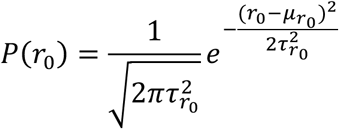

- Prior on 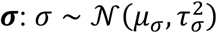, where *μ*_σ_ and 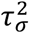 are the mean and variance of the prior on *σ*.

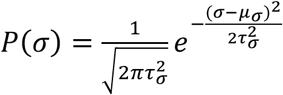

The goal is to find the posterior distribution *P*(*r*_0_, *σ* ∣ *r*_1_, *r*_2_, …, *r_n_*), which is proportional to the product of the likelihood and the prior:

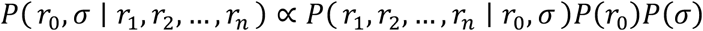

Substituting the likelihood and priors:

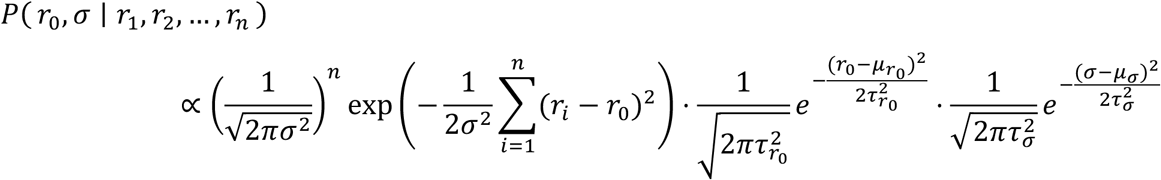

Combining the exponentials:

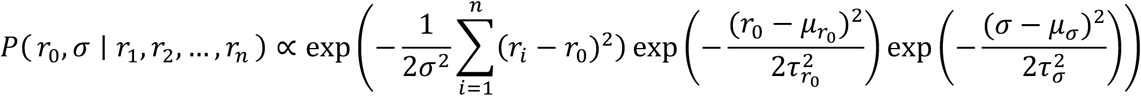

This can be expressed more compactly as:

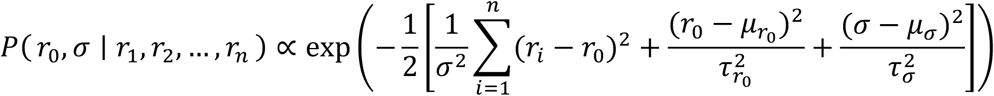

#### MCMC Framework (Metropolis-Hastings Algorithm)

The Metropolis-Hastings algorithm involves iteratively proposing new values for model parameters based on a predefined distribution and calculating an acceptance ratio that compares the likelihood of the new parameters against the current ones. If the new parameters yield a higher probability or meet a random acceptance criterion, they are accepted; otherwise, the current parameters are retained. This process is repeated for a set number of iterations, with each accepted parameter set stored for subsequent analysis. The method ensures exploration of the parameter space while gradually converging on the most probable values based on the observed data.

### Statistical Analysis

To evaluate differences between groups for each metric, we conducted statistical testing in a two-step process. First, pairwise comparisons were performed using the Mann-Whitney U test (Mann and Whitney 1947), a rank-based non-parametric method for comparing two independent groups.

This test evaluates whether one group tends to have higher or lower values than the other, without assuming normality or equal variances. We compared all groups to a reference group, Complete Spatial Randomness (CSR), for each metric. To control for multiple comparisons and reduce the likelihood of false positives, we applied the Benjamini-Hochberg False Discovery Rate (FDR) correction (Benjamini and Hochberg 1995) to the raw p-values. For each comparison, the Mann-Whitney U test results included the test statistic, raw p-value, corrected p-value, and a significance status (whether the corrected p-value was below 0.05). Significant results were bolded for presentation in the final dataset. The corrected p-values were used to determine statistical significance, with values below 0.05 considered indicative of significant differences.

### Metrics for Comparing Distributions

We employed three metrics—Wasserstein Distance, Normalized Mean Squared Error (MSE), and the Kolmogorov-Smirnov (KS) Statistic—to compare the similarity between real and synthetic data distributions. These metrics were calculated for multiple methods and across two conditions: scrambled (control) and siNCAPH2 (treated) and for the eight cell line distributions. These metrics assess the level of agreement between two probability density functions (PDFs), enabling us to quantify the fidelity of synthetic data in capturing the characteristics of experimental data.

The Wasserstein Distance (WD), also known as the Earth Mover’s Distance, measures the minimum “cost” of transforming one distribution into another. It is defined for one-dimensional distributions as:

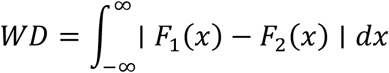

where *F*_1_(*x*) and *F*_2_(*x*) are the cumulative distribution functions (CDFs) of the two datasets. Lower Wasserstein distances indicate greater similarity.

The Normalized Mean Squared Error (MSE) quantifies the discrepancy between the real and synthetic PDFs by normalizing the MSE by the variance of the real PDF. It is given by:

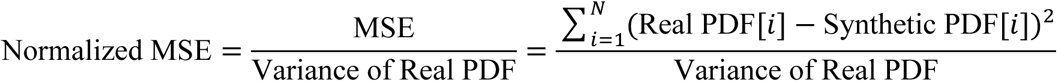

A lower normalized MSE indicates higher similarity, with values close to zero reflecting strong agreement.

Finally, the Kolmogorov-Smirnov (KS) Statistic compares the maximum difference between the CDFs of two distributions, defined as:

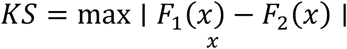

where *F*_1_(*x*) and *F*_2_(*x*) are the cumulative distribution functions (CDFs) of the two datasets. Smaller KS values indicate closer alignment between the distributions. Together, these metrics provide a comprehensive assessment of the similarity between real and synthetic data across various methods and experimental conditions.

## Results

### Simulation of Centromeric Spot Patterns Using Different Spatial Distribution Models

To test and benchmark centromere clustering metrics we first generated synthetic image sets representing a range of spatial patterns of centromeres, including uniform distributions, clustered arrangements, and dispersed configurations, to replicate centromere arrangements observed in experimental data (Kiskowski, Hancock, and Kenworthy 2009). We used 9 different spatial distribution models to mimic the centromeric localization observed in HCT116-Cas9 colorectal cancer cells stained for the CENP-Centromere protein CENP-C (Figure 1), which stably binds to centromeres in interphase nuclei (Fachinetti et al. 2015). The models used for synthetic data generation included: a Complete Spatial Randomness (CSR) that draws samples from a 2D uniform distribution (Figure 1A), a Poisson-Disk Sampling (PDS) process that sets a minimum distance threshold between two neighboring spots (Figure 1B), Uniformly distributed with Two Adjacent spots (UTA) (Figure 1C) or Uniformly distributed with Three Adjacent spots (UTHA) (Figure 1D) with adjacent spots mimicking the formation of small clusters, Single Two-Dimensional Gaussian (S2DG) (Figure 1E), Two Two-Dimensional Gaussian (T2DG) (Figure 1F), or Three Two-Dimensional Gaussian (TH2DG) (Figure 1G) to represent clustering in large nuclear regions, and either Two-Dimensional Gaussian with a Nuclear Body (2DGNB) (Figure 1H) or Two Two-Dimensional Gaussian with Two Nuclear Bodies (T2DGTNB) (Figure 1I) with central exclusion to mirror centromeres localized in proximity of large nuclear bodies, such as nucleoli. For each of these spot patterns we generated 100,000 synthetic images of nuclei, assuming nucleus shape to be circular with a diameter of 10 micrometers, based on measurements of the mean nucleus area obtained from experimental image datasets of HCT116-Cas9 cells (Keikhosravi et al. 2024).

### Benchmarking of Clustering Metrics on Synthetic Patterns

These synthetic spot distribution datasets offer a controlled environment for assessing how well different clustering metrics can detect specific clustering patterns. Accordingly, we used them to benchmark six clustering metrics (Figure 2) to measure local and global differences in point clustering patterns (Kiskowski, Hancock, and Kenworthy 2009; Savulescu et al. 2021; Shizuka and Farine 2016; Norton et al. 2018). In particular, we focused on how sensitive each metric was to detecting differences in clustering patterns between the synthetic data simulated by a Complete State of Randomness (CSR) that draws samples from a 2D uniform distribution, and by each of the other distributions that represent either increased degrees and patterns of spot clustering, or to detect centromere dispersion for the Poisson-Disk Sampling (PDS) (Figure 2, see Materials and Methods for details). When applied to synthetic centromere spot patterns, the clustering metrics showed clear differences in their sensitivity to detect different centromere clustering patterns (Figure 2A).

**Figure 2.**
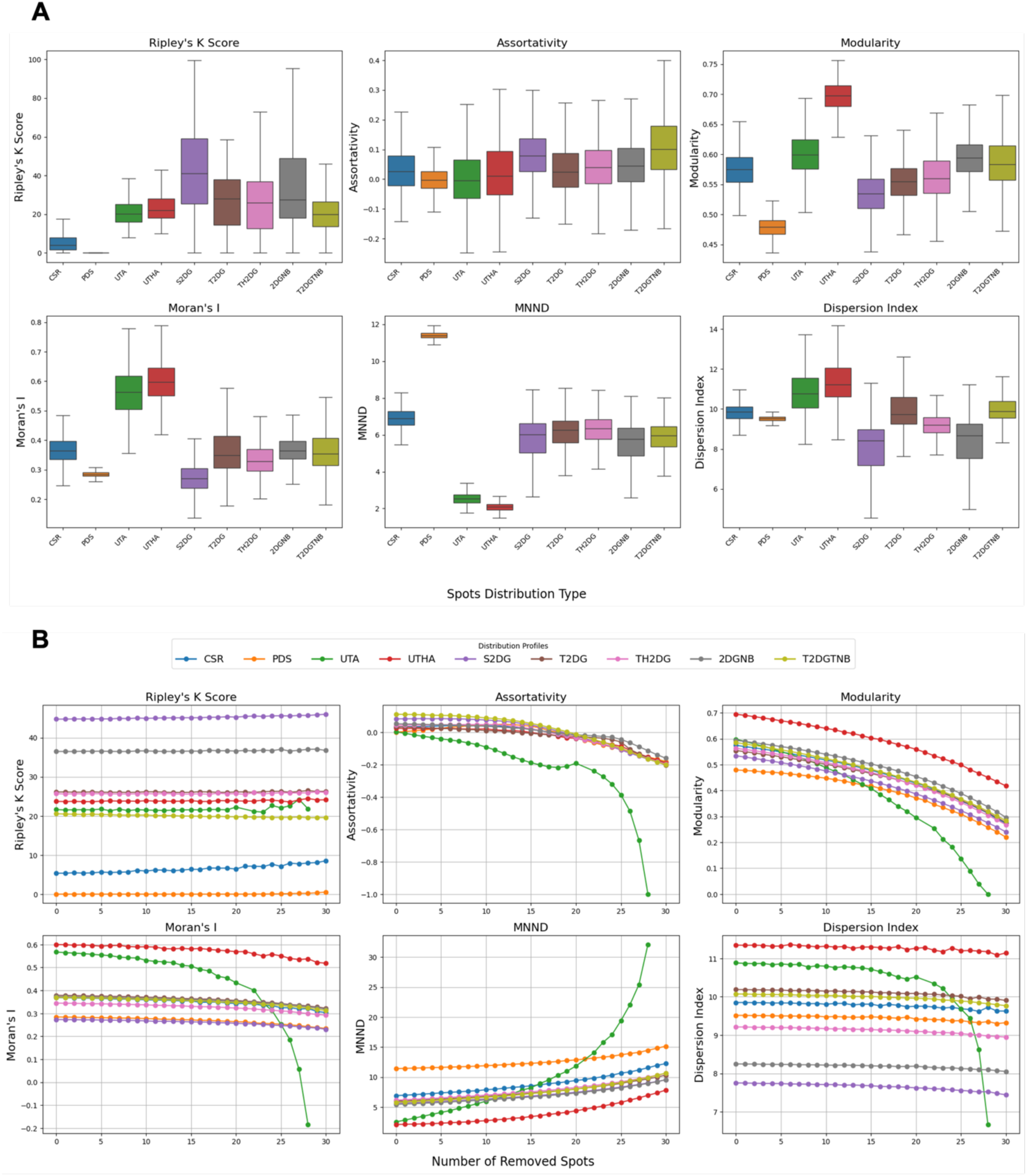
Benchmarking clustering metrics for detecting and characterizing synthetic centromere clustering patterns. (A) Sensitivity of the six clustering metrics was tested across synthetic spot distributions. Each boxplot displays the variability of metric values for the nine spatial distribution models, with statistical testing confirming significant differences between CSR and other distributions for most metrics. The box represents the interquartile range (IQR), showing the middle 50% of the data, with the line inside the box indicating the median value. The whiskers extend from the box to the smallest and largest values within 1.5 times the IQR from the lower and upper quartiles. (B) Robustness of clustering metrics to changes in spot number was evaluated by progressively removing a random number of spots from synthetic nuclei.

The Ripley’s K Score (Ripley 1976; Kiskowski, Hancock, and Kenworthy 2009), had a value of zero for PDS and very small values for uniformly distributed spots, most likely due to the limited area and spots sampling size. Pairwise Mann-Whitney U tests, corrected using the Benjamini-Hochberg False Discovery Rate (BH-FDR) method, showed a statistically significant difference when using Ripley’s K Score between CSR and all other spot generation methods (p < 0.05) (Figure 2A and Supplementary Table 2), making it a robust metric for measuring centromere clustering. Cohen’s D values, also indicated a substantial difference between the same comparison groups (Supplementary Figure 1). The Assortativity Index is a measure of connectedness of nodes with similar degrees in a graph (Shizuka and Farine 2016). For this metric, statistical testing showed a significant difference between CSR and all other spot generation methods (p < 0.05), except for T2DG (Figure 2A). However, the absolute Cohen’s D values for this metric showed only small (D < 0.2) to medium (0.5 < D <0.8) (Supplementary Figure 1) differences between clustering patterns and CSR negative control, limiting its use for detection of changes in centromere clustering patterns. The Modularity Index, a measure of connection density within clusters compared to connections between clusters in a graph, reflects how well the graph divides into modules. The Modularity Index analysis showed a statistically significant difference between CSR and all other spot generation methods (p < 0.05) (Figure 2A and Supplementary Table 2). However, this metric only showed a substantial difference between CSR and three other spot generation methods: PDS, UTHA and S2DG, while the rest were either modest or small (Supplementary Figure 1). Moran’s I is a measure of spatial autocorrelation (Tiefelsdorf and Boots 1995). Statistical testing yielded a significant difference between CSR and all other spot generation methods (p < 0.05), except for 2DGNB (Supplementary Table 2). This metric, however, only showed a substantial difference between CSR and three other spot generation methods: PDS, UTA, UTHA and S2DG, while the rest are either modest or small (Supplementary Figure 1). Mean Nearest Neighbor Distance (MNND) (Clark and Evans 1954), which measures the average distance to the nearest neighboring spot, showed a significant difference between CSR and all other spot generation methods (p<<0.05) (Supplementary Table 2). Cohen’s D values of above 0.8 also confirms a substantial difference between the same comparison groups, except TH2DG, which was only modest (0.5 < D <0.8) (Supplementary Figure 1). Dispersion Index (Myers 1978), calculated as the variance over mean of the pairwise distance distribution, shows a significant difference between CSR and all other spot generation methods (p<<0.05) (Supplementary Table 2). Cohen’s D values of above 0.8 also confirmed substantial difference between the same comparison groups, except for T2DG and T2DGTNB which are small (0.2 < D <0.5) (Supplementary Figure 1).

These results suggest that while some different metrics such as MNND and Dispersion Index perform well in separating CSR from other distributions, only the Ripley’s K Score showed robust and consistent differences between CSR and both clustering and dispersion patterns. We conclude that Ripley’s K Score is the most suitable clustering metric to detect a wide range of clustering pattern as would for example be seen in unbiased screening approaches (Supplementary Figure 1 and Supplementary Table 2).

### Measuring Clustering Metrics Robustness to Synthetic Changes in Spot Number per Nucleus

Since the human HCT116 colon cancer cell line is pseudodiploid (Rajput et al. 2008) and the expected number of detected centromeres in these cells is 46. However, we, that the median number of detected CENP-C-labelled centromere spots per nucleus in HCT116-Cas9 cells was only 32 (Keikhosravi et al. 2024). This discrepancy is possibly due to the limited optical resolution of our diffraction limited microscopes, which cannot resolve multiple centromeres located very closely to each other in maximal projections of image z-stacks (see Materials and Methods). Given the lower number of detected centromere signals in experimental data compared to simulated spot distributions, we sought to determine how sensitive the clustering metrics were to changes in CENP-C spot number. To address this question, we re-tested all clustering metrics on the synthetic spot image datasets but removed at random a variable number of spots (from 1 to 30) from each nucleus (Figure 2B). The average value for each metric was then determined for each number of spots removed from the initial 46 spots, and the percent change was calculated to quantify the relative variability of each metric across all distributions compared to the value with 46 spots (Figure 2B; Supplementary Table 3). Once again, Ripley’s K Score proved to be the most robust metrics. Both for Ripley’s K Score and for the Dispersion Index most percent change values were below 10%, indicating robustness of these metrics to the number of detected spots. On the contrary, Moran’s I and Modularity, showed a moderate variation with the number of spots (below 60%), while Assortativity and MNND showed high sensitivity for most of spot generation models (Supplementary Table 3).

Altogether, the results of these benchmark tests on synthetic images indicate that the Ripley’s K Score, and to a lesser extent, the Dispersion Index, which are both directly calculated from the statistical properties of spot distance distributions in a cell, are the most robust to spot number variability (Figure 2B). The Ripley’s K Score also showed the best performance in separating different distributions based on Cohen’s D values (Supplementary Figure 1). Other metrics, however, may still be used for finding specific patterns (such as Modularity and MNND for adjacent spots), albeit with less reliability.

### Validation of Clustering Metrics Using Experimental Data

Having benchmarked the different clustering metrics on synthetic images, we next tested all metrics on HTI images of cells stained with an antibody against the centromere protein CENP-C in HCT116-Cas9 cells (Figure 3 A). Since one of the major applications of a quantitative clustering metric would be its use in detecting changes in clustering patterns, for example in imaging-based optical screens, we also sought to determine which clustering metric produced the largest difference between the normal distribution of centromeres in HCT116-Cas9 cells and HCT116-Cas9 cells transfected with an siRNA oligo targeting the *NCAPH2* gene, which encodes for a subunit of the condensin II complex, and whose knock-down is known to lead to increased centromeric clustering (Figure 3A) (Keikhosravi et al. 2024; Hoencamp et al. 2021). The results of this experiment revealed varying abilities of the different clustering metrics to capture this known biological effect (Figure 3B).

**Figure 3.**
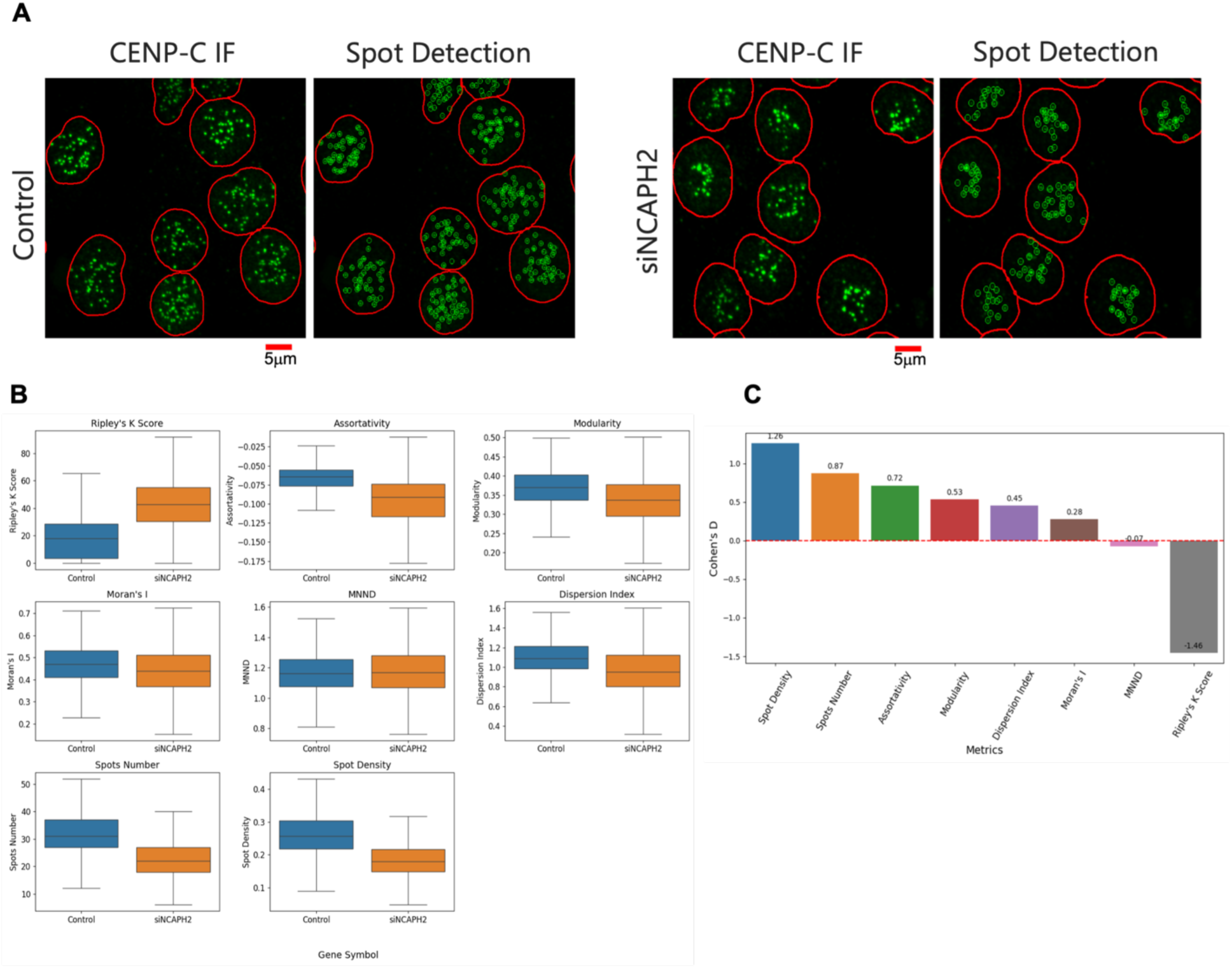
Analysis of centromere clustering in control and NCAPH2-depleted cells. (A) Representative immunofluorescence images showing CENP-C spot detection (green) in HCT116-Cas9 cells transfected with siScramble (control) or siNCAPH2. The nuclear outline is indicated in red. Scale bar: 5μm. (B) Comparison of different clustering metrics to quantify differences between control and NCAPH2-depleted cells. The box represents the interquartile range (IQR), showing the middle 50% of the data, with the line inside the box indicating the median value. The whiskers extend from the box to the smallest and largest values within 1.5 times the IQR from the lower and upper quartiles. (C) Cohen’s D analysis showing the relative sensitivity of various clustering metrics for separating siScrambled (control) and siNCAPH2 in HCT116 cells, with Ripley’s K demonstrating the highest absolute value, followed by Spot Density, Spot Number, Modularity, and Assortativity.

While each of the clustering metrics could detect differences in centromere distribution at the single cell level (p < 0.05), the Ripley’s K clustering score produced the largest difference in centromere clustering in HCT116-Cas9 cells between negative control and NCAPH2 knock down cells, followed by Modularity and Assortativity (Figure 3B). Similarly, when we also calculated and plotted values for the CENP-C spot number and density on a per cell basis, we observed large differences upon NCAPH2 knockdown, likely due to increased clustering of individual centromeres into larger spot aggregates that cannot be resolved with diffraction limited microscopy. Accordingly, the Cohen’s D value for all these metrics indicated that the Ripley’s K clustering Score was again the most sensitive metric followed by Spot Density, Spot Number, Modularity and Assortativity (Figure 3C). Using the Dispersion Index, Moran’s I, and MNND for the same image set led to much lower Cohen’s D values (Figure 3 C). The results of these experiments indicate that use of Ripley’s K is the most sensitive metric to detect changes in overall clustering of centromeres in cells.

### Modelling the Spatial Distribution of Centromeres in Cells

As a complementary approach to our identification of the most sensitive metrics for analysis of centromere distributions, we asked whether we could predict the observed distribution of centromere patterns by a modelling approach. Towards that goal, we first sought to establish the overall spatial distribution of centromeres in HCT116-Cas9 cells. Accordingly, we overlaid all the spots detected in HCT116-Cas9 cells by shifting all spots within each cell so that the nuclei centers were located at (0,0) position (Figure 4). The histogram of the standardized distribution of the spots in 2D space revealed a doughnut-like shape with the center at (0,0) (Figure 4A). A line plot of the 2D histogram of standardized spot locations confirmed low spot density near the center of the nucleus and a higher density in the mid-region between the center and the edge of the nucleus (Figure 4B). It is likely that the low density of the spots at the center is caused by the well-established presence of major nuclear compartments such as the nucleolus in the interior of the nucleus (Naughton et al. 2022; Kumar et al. 2024; Rodrigues et al. 2023).

**Figure 4.**
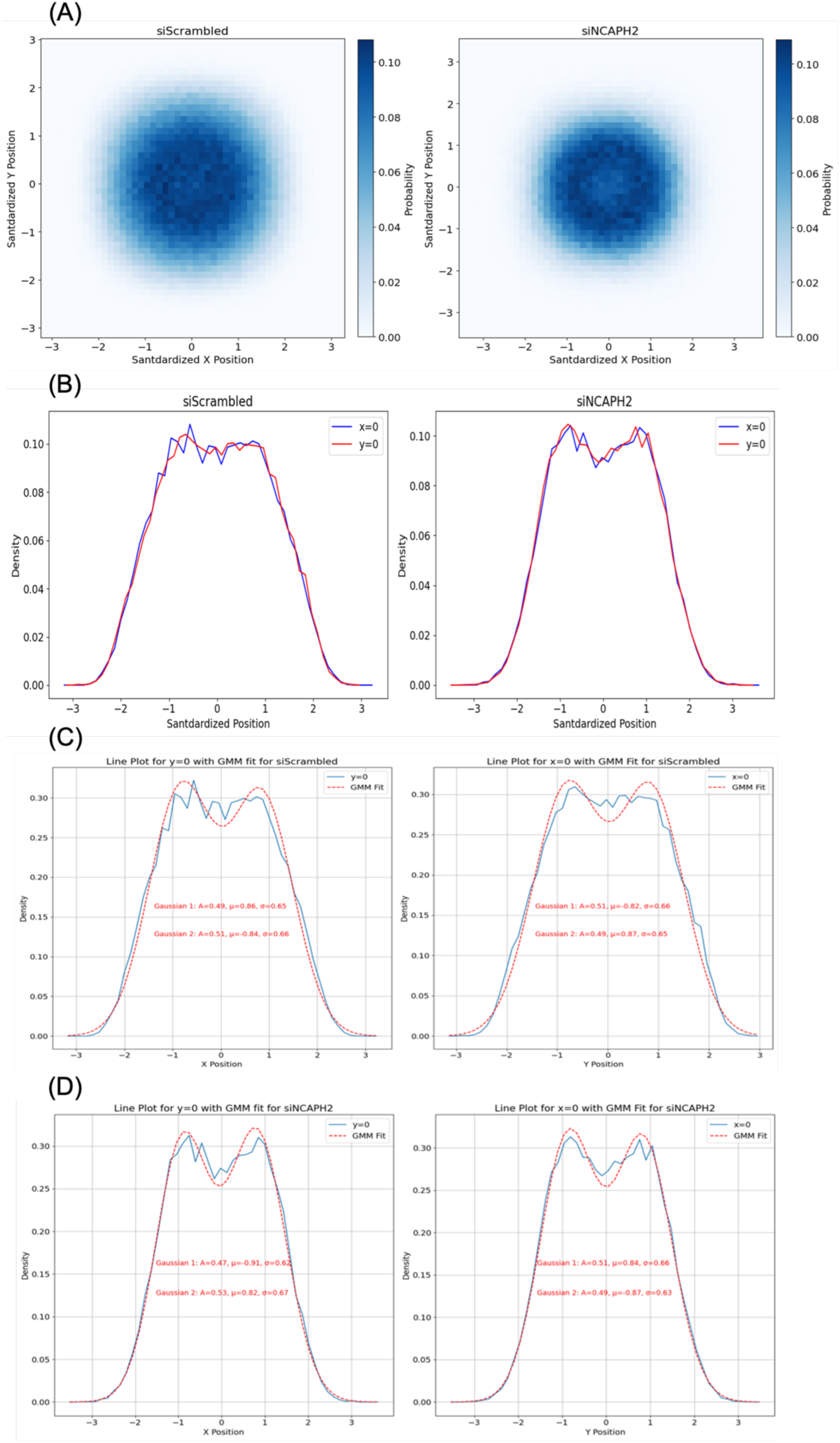
Spatial distribution of standardized centromere locations in control and NCAPH2-depleted cells. (A) 2D histogram showing the distribution of CENP-C spots relative to the nuclear center (0,0), revealing a doughnut-shaped pattern in both conditions. (B) Line plot analysis at x=0 and y=0 demonstrating lower spot density at the nuclear center and higher density between center and nuclear edge. (C, D) Gaussian mixture modeling of spot distribution along x=0 and y=0 axes, indicating radially symmetric centromere organization with potential alignment artifacts affecting the observed central low-density region.

The distribution in both X and Y directions could be accurately modeled using a 2-mode one-dimensional Gaussian mixture (Figure 4C, D). The closeness of the Gaussian parameters for fitted line plots in both directions (Pearson Correlation Coefficient > 0.99), suggested that a radial Gaussian distribution that is uniformly distributed in all directions reflects well the cellular distribution of centromeres. The same properties of a doughnut-like distribution and close fit to a radial Gaussian distribution were found upon knockdown of NCAPH2 (Figure 4E-H).

### Centromere Localization Modeling using Parametric Distributions

To assess the centromere localization in 2D using parametric distributions based on observations in Figure 4, we fit several spatial distribution models to the spot locations acquired using imaging data. Then, to measure the accuracy of the simulations, we first simulated spot positions overlaid on nuclei masks resulting from the DAPI channel segmentation using the parameters obtained from the models fit from the microscopy images. We then compared the simulated pairwise spot-to-spot distance distributions and the spot radial distance distributions with the ones obtained from the microscopy images (Keikhosravi et al. 2024).

We used several spot distributions for modeling of the experimental imaging data: 1) Complete State of Randomness or uniformly distributed spots in 2D (Model 1; M1) with no preferential localization; 2) Nucleus shaped Gaussian (M2), which represents a radially distributed pattern with parameters constrained by the nucleus shape; 3) Radial Gaussian distribution in 2D space with uniformly distributed spots for all angles. For this method we extracted the model parameters by fitting to the analytically calculated CDF of pairwise distances to the same function from real data (M3; see Materials and Methods for details); 4) Radially shifted Gaussian, assuming uniform distribution across 360 degrees. The model parameters were calculated in an iterative approach using a Bayesian framework that used the spot coordinates from real data (∼46 spot coordinates per cell) (M4; see Materials and Methods for details); 5) Radially shifted Gaussian distribution using spot distances, assuming uniform distribution across 360 degrees. The model parameters were calculated in an iterative approach using a Bayesian framework that used the distribution of pairwise distances of spots from real data (∼1000 pairwise distances per cell) (M5; see Materials and Methods for details).

By comparing the simulated (M1–M5) spot localizations with the observed CENPC-C spot positions measured in images of HCT116-Cas9 cells (M0), we observed varying degrees of similarity for both pairwise and radial distance distributions (Figure 5A). The pairwise distance distribution generated by the M4 model most closely fitted the experimental data, with its density curve aligning well at the peak and along the tail regions, indicating its high accuracy in capturing the underlying spatial organization of centromeres (Figure 5A). M3 and M5 also performed significantly better than M1 and M2, but their peak was slightly less aligned with experimental data compared to M4. These observations were quantified using three established distribution similarity metrics: Wasserstein Distance, Normalized Mean Squared Error (MSE), and KS Statistic, which quantify different properties of a distribution (see Materials and Methods for details) (Figure 5B). These analyses indicate a better fit to the observed distributions using the M4 model followed by M3 and M5, with most of its distribution similarity metrics below 10% (Figure 5B).

**Figure 5.**
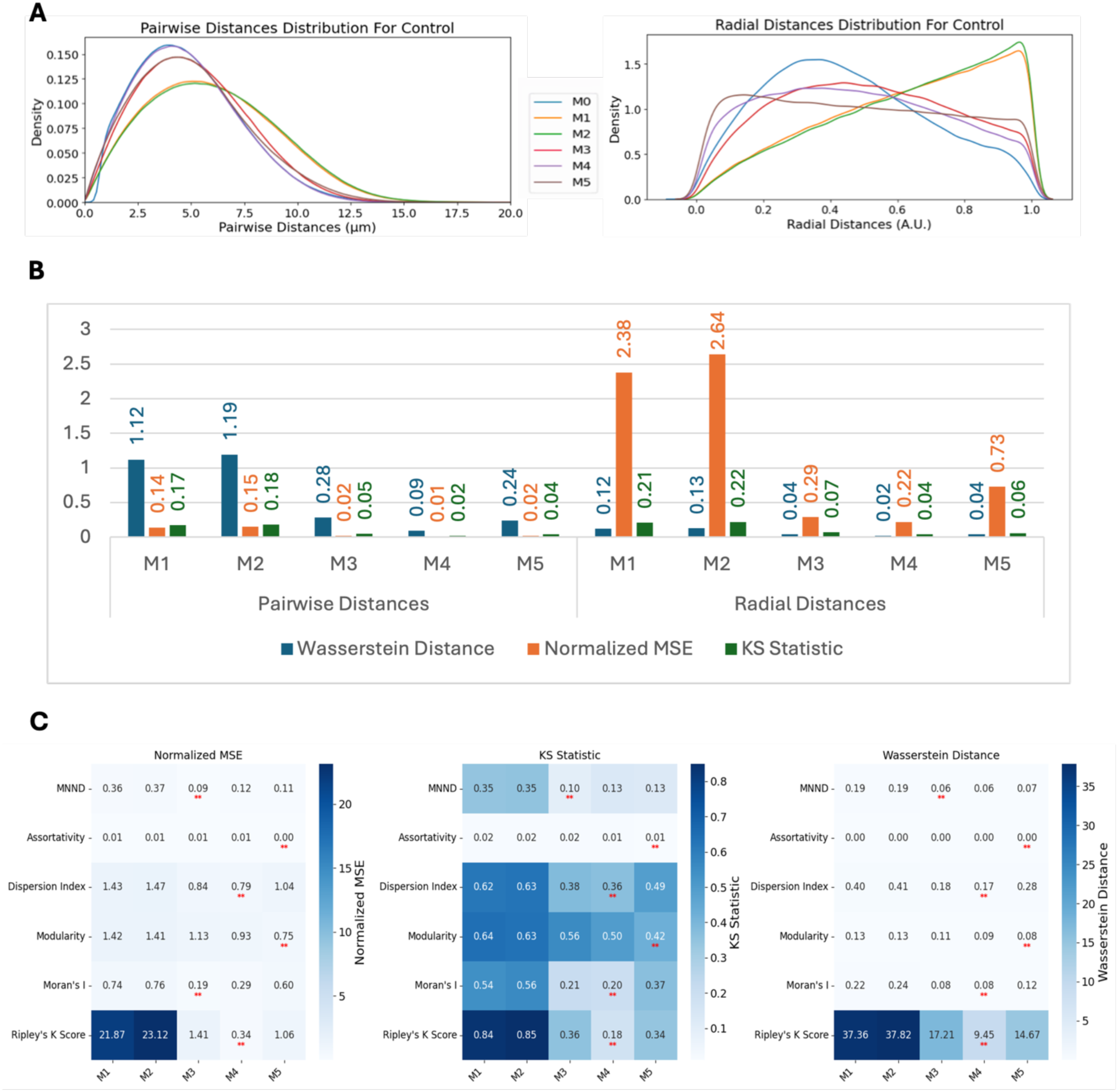
Comparative analysis of centromere spatial distribution models. (A) Pairwise and radial distance distributions comparing experimental data (M0) with five different synthetic models (M1-M5) in HCT116-Cas9 cells. (B) Quantitative assessment of model performance using Wasserstein Distance, Normalized MSE, and KS Statistic, showing superior performance of M4. (C) Heatmap analysis of clustering metrics across different models, demonstrating that Bayesian-based models (M4 and M5) achieve the lowest Normalized MSE values for most metrics.

### Spot clustering metrics comparison for generated spots

We also compared the spot clustering metrics for spots generated using various spot modeling methods (Figure 5 C). The Bayesian-based models (M4 and M5) consistently achieved the lowest Normalized MSE values across most metrics. Metrics such as Ripley K Score, Modularity, and Dispersion Index showed significant improvement with M4 and M5 compared to simpler models like M1 and M2, highlighting their ability to replicate global spatial clustering behavior. Although M4 and M5 did not outperform other methods for MNND, the differences between these models and the best-performing methods were very minimal. The values for the Assortativity Index in most cases were below 0.005 and displayed as zero. This is in line with our previous observation when comparing different metrics for varying number of spots (Figure 2B), indicating that the Assortativity is less effective in discriminating different spot distributions.

We next asked whether spot localizations simulated using models M1 – M5 can approximate the localization of CENP-C spots when centromeric spatial patterns are perturbed s experimentally, such as when NCAPH2 is knocked down (siNCAPH2) in HCT116-Cas9 cells, which leads to increased clustering of centromeres (Ono et al. 2017; Wallace et al. 2019; Martin et al. 2016). Upon analysis of the siNCAPH2 dataset, M4 still remained the best model followed by M3 and M5 (Figure. 6A). While quantification of these observations using Wasserstein Distance, Normalized MSE, and KS Statistic further confirmed M4’s superior performance followed by M3 and M5 (Figure 6B), the similarity between modeled distributions and real data distribution of pairwise and radial distance distributions was decreased compared to the control HCT116-Cas9 cells (Figure 5B). This could be in part due to smaller numbers of spots detected in siNCAPH2 treated cells, which lowers the accuracy of the model fitting, as well as due to changes in cellular architecture due to NCAPH2 loss.

**Figure 6.**
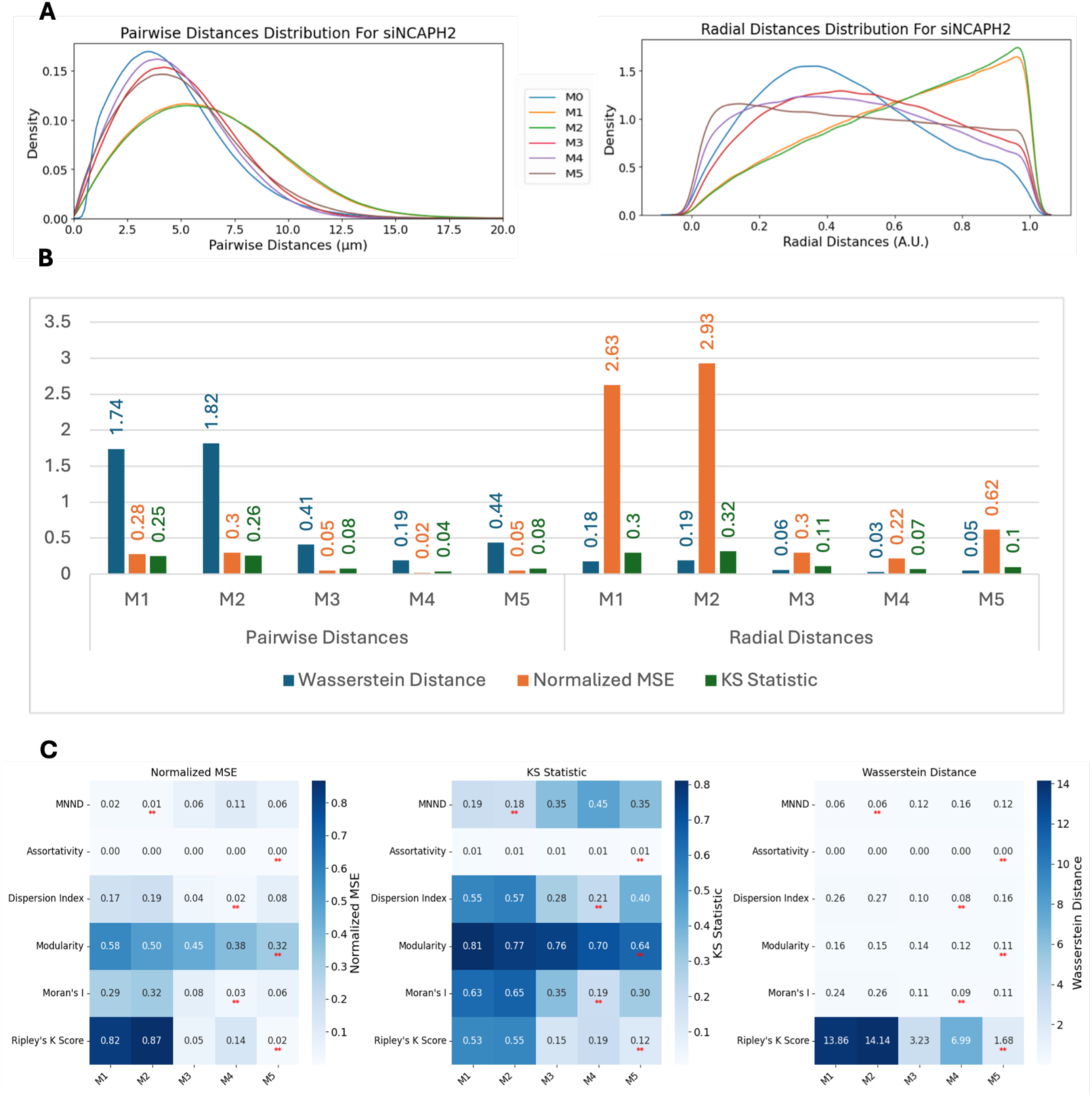
Model performance analysis in NCAPH2-depleted cells. (A) Pairwise and radial distance distribution comparisons between experimental data and synthetic models (M1-M5) in siNCAPH2-treated cells. (B) Distribution similarity metrics showing M4 model maintains best performance but with reduced accuracy compared to control cells. (C) Clustering metric analysis demonstrating Bayesian models (M4, M5) achieve lowest similarity values for most metrics, with slight variations in MNND performance.

Comparing spot clustering metrics for spots generated using the various spot modeling methods with the same metrics calculated using the experimental data for siNCAPH2 cells (Figure 6C) showed a similar trend as in control cells (Figure 5C). Similar to control cells, in siNCAPH2 datasets, M4 and M5 achieved the lowest values for similarity metrics such as Ripley K Score, Modularity, Moran’s I and Dispersion Index, confirming their robustness in capturing centromere clustering patterns.

### Evaluation of Generative Models for CENP-C Spot Localization Patterns in Multiple Human Cell Lines

To finally ask whether these quantification methods are generally applicable to multiple cell types, we tested them against imaging dataset from a diverse set of eight human cell types and tissues, ranging from colon cancer cells and skin cells to induced pluripotent stem cells which show an exceptionally high level of clustering (Rodrigues et al. 2023). Similar to HCT116-Cas9 cells, we observed the same pattern of doughnut-like localization by overlaying the nucleus centered spot locations in all eight cell lines (Figure S2A). All cell lines showed a similar Gaussian shaped distribution of spot densities (Figure S2B). Visual inspection of the pairwise distance distributions revealed that model-based distributions can reliably reproduce the distribution of distances with high accuracy (Figure 7A). While the models were not fitted based on radial distances, they could capture the underlying radial positioning (Figure 7B).

**Figure 7.**
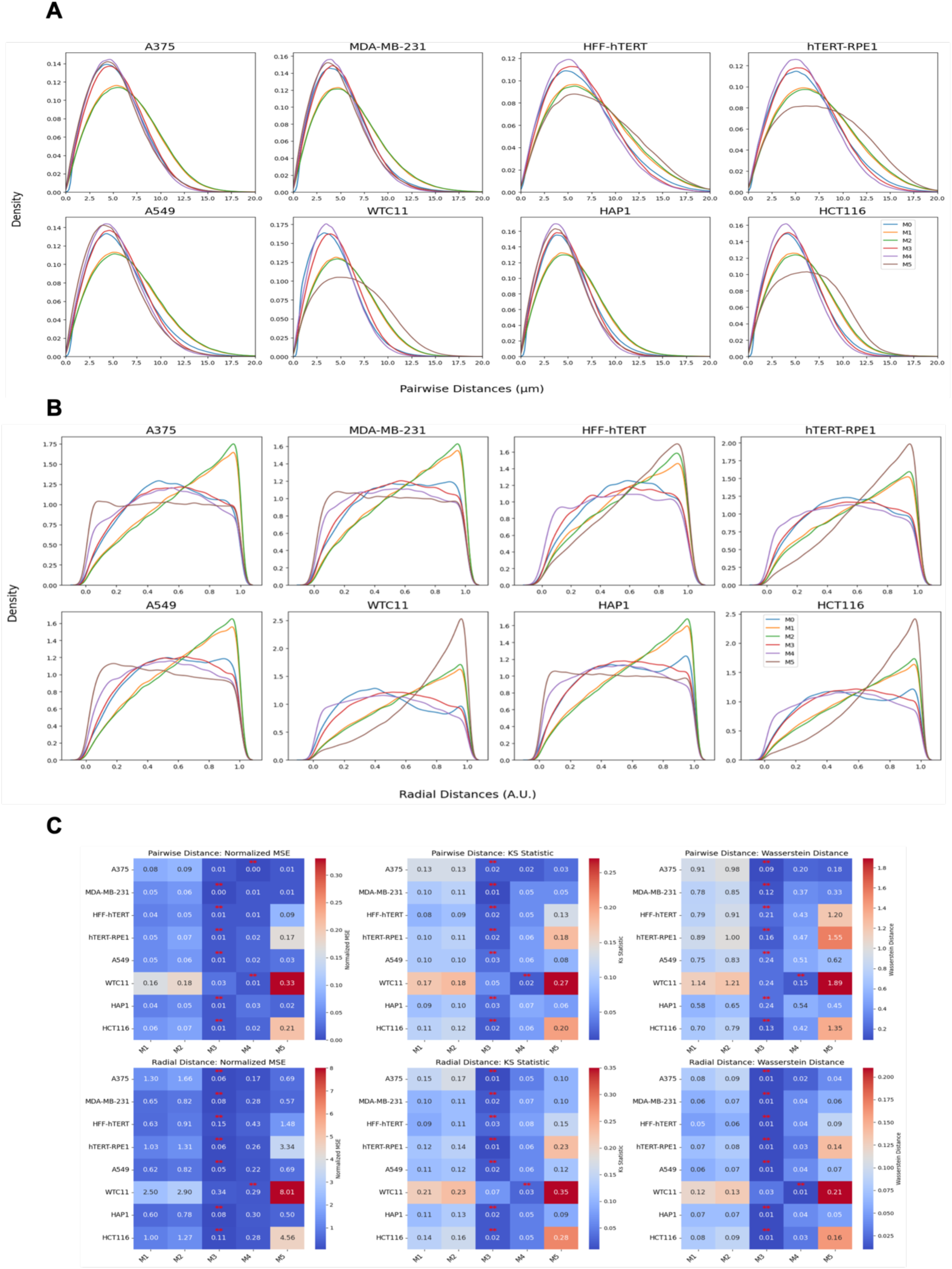
Centromere spatial organization analysis across cell lines including HCT116 (colon), A375 (melanoma), MDA-MB-231 (breast), HFF-hTERT (fibroblast), hTERT-RPE1 (retinal), A549 (lung), HAP1 (myeloid), and WTC11 (embryonic stem cells). (A) Pairwise distance distributions comparing experimental data with model predictions. (B) Radial distance distributions showing models’ ability to capture centromere positioning. (C) Quantitative comparison of model performance using Wasserstein Distance, Normalized MSE, and KS Statistic, demonstrating M3 as the best-performing model across most cell lines, with M4 as second-best.

We did notice a clear difference in the performance of the computational models in reproducing centromere distributions in various cell types (Figure 7C). For pairwise distances, except the WTC11 embryonic stem cell line, the M3 model consistently achieved the lowest Wasserstein Distance, Normalized MSE, and KS Statistic across most cell lines, indicating that it is the most effective at predicting the observed pairwise distance distributions of centromeres. Similarly, M3 also outperformed other methods when radial distribution was analyzed. M4 followed closely as the second-best performing model, followed by M5, M1 and M2. This trend was observed across most cell lines. M5 exhibited a noticeably weaker predictive performance compared to the other methods. This could potentially be attributed to the small number of cells available which is not sufficient for the more complex M5 model that uses a larger input set (>1000 pairwise distances) to fully capture the spatial organization of centromeres.

## Discussion

Understanding the spatial organization of cellular structures is a fundamental question in modern cell biology. Centromeres are prominent cellular features of each chromosome and elucidation of their distribution in the human cell nucleus is crucial for deciphering chromosomal behavior and nuclear architecture in both normal and disease cells. To this end, we have generated a systematic framework and specific measurement tools to quantitatively assess the cellular distribution of human centromeres. These new methods will be useful in the investigation of the mechanisms involved in establishing and maintaining centromere localization patterns and their functional implications.

We used this framework to assess multiple centromere spatial distribution types, clustering metrics, and spot generation models. By simulating diverse spatial patterns—including uniform distributions, single Gaussian clusters, multi-modal Gaussian mixtures, and perturbed adjacent spots—these models allow precise comparison and identification of the metrics most suited for specific biological questions. This benchmarking process ensures that the chosen metric aligns with the spatial characteristics of interest, enhancing the rigor and relevance of quantitative assessments.

Based on our analysis, the Ripley’s K Score emerged as the most sensitive and versatile metric for measurement of global centromere clustering, with minimal variation across different numbers of detected spots. This property makes it a suitable metric for studying changes in centromere clustering upon experimental perturbation for example after elimination of the condensin protein NCAPH2. Both sensitivity and robustness of this metrics are mainly due to the fact that it is calculated using the density normalized CDF of all spot distances, irrespective of local features of their distribution. Although Dispersion Index is also using the first and second order statistics of the same distribution, but the compression of the distribution into only two parameters, reduces its effectiveness in separating various spot distributions compared to Riley’s K Score. One limitation of Ripley’s K Score may be its ability to detect local clusters dispersed within the nucleus because it is calculated using the overall distance distribution of spots.

To better understand the nature of cellular centromere distribution, we used computational modeling to replicate the localization and clustering behavior of centromeres. Specifically, we applied Bayesian estimation methods to fit a radially shifted Gaussian distribution using three different approaches. The first approach directly fit the cumulative distribution function (CDF) of pairwise distances between centromeres. The second approach used spot coordinates (approximately 46 spots per cell), while the third relied on pairwise distances (around 1,035 values per cell). Although all these models are based on radial Gaussian distributions, the third approach, with its larger number of input values, requires a higher sample size to achieve robust and accurate parameter fitting. This highlights the trade-off between input complexity and the need for larger datasets when modeling spatial distributions.

One of our findings based on mapping the location of several thousand centromeres by imaging, is that centromeres are distributed in a doughnut-like distribution within the cell nucleus. This pattern was evident when mapping individual centromeres across cell populations and was reproduced in our modeling approaches. This finding aligns well with previous studies on the radial organization of nuclear structures, such as observation of lower density of chromatin and chromosomes in the nuclear interior (Kreth et al. 2004; M. Cremer et al. 2001). These studies highlight the influence of biological constraints, such as gene density and chromatin architecture, as key drivers of spatial nuclear organization.

The analytical tools we have generated have practical applications. These analysis methods can be used to quantitatively describe changes in centromere distribution in response to a specific experimental perturbation, for example, loss of the cohesin component NCAPH2 as shown here, or during physiological or pathological processes such as differentiation, development and in disease such as cancer. We also anticipate that the methods are applicable to analysis of tissue samples, provided nuclei can be segmented accurately. Maybe more importantly, our analysis approaches will now enable the execution of large scale functional genomic screen, such as CRISPR-screens, which will allow the identification of entirely novel modulators of centromere localization in an unbiased fashion.

The combination of advanced imaging techniques with robust computational analysis, as demonstrated here, are not limited to centromeres but can also be extended to study the spatial organization of other cellular structures. For instance, nuclear bodies such as nucleoli, Cajal bodies, and PML nuclear bodies exhibit distinct spatial patterns influenced by chromatin interactions and nuclear architecture (Dundr and Misteli 2010). Probabilistic and spatial modeling approaches, like those described here, have already been applied to chromosomal territories and transcription factories, revealing the roles of gene density and transcriptional activity in nuclear organization (M. Cremer et al. 2001; Kreth et al. 2004; Fraser and Bickmore 2007). Additionally, approaches using spatial modeling or clustering metrics can be tailored to study other biological structures, such as cytoplasmic structures like stress granules (Wheeler et al. 2016), focal adhesions (Kanchanawong et al. 2010) or mitochondrial networks (Picard et al. 2013), to better understand their distribution and functional implications. By adopting and adapting these methods, the principles underlying spatial organization and its impact on cellular processes across a wide range of subcellular structures can be explored using quantitative measures.

Taken together, this study presents a framework for characterizing the cellular distribution of centromeres using a combination of spatial metrics and computational modeling. By systematically benchmarking metrics with synthetic spot generation models, we demonstrate the capability of combining data-driven analysis with computational tools to capture centromere clustering patterns with potential application to other cellular structures.

## Code Availability

All the code used for modeling and generating the results and plots in this manuscript can be found here: https://github.com/CBIIT/centromere_clustering_analysis.

## Acknowledgements

We would like to thank Thomas Gonatopoulos-Pournatzis for sharing several of the cell lines used in this study. This work utilized the computational resources of the NIH HPC Biowulf cluster (https://hpc.nih.gov). Work in the Misteli Lab is supported by the Intramural Research Program of the NIH, NCI, Center for Cancer Research through grant 1-ZIA-BC010309-25. HiTIF was supported by the Intramural Research Program of the NIH, NCI, Center for Cancer Research through grant ZIC BC 011567.

---

**Supplementary Figure 1.**
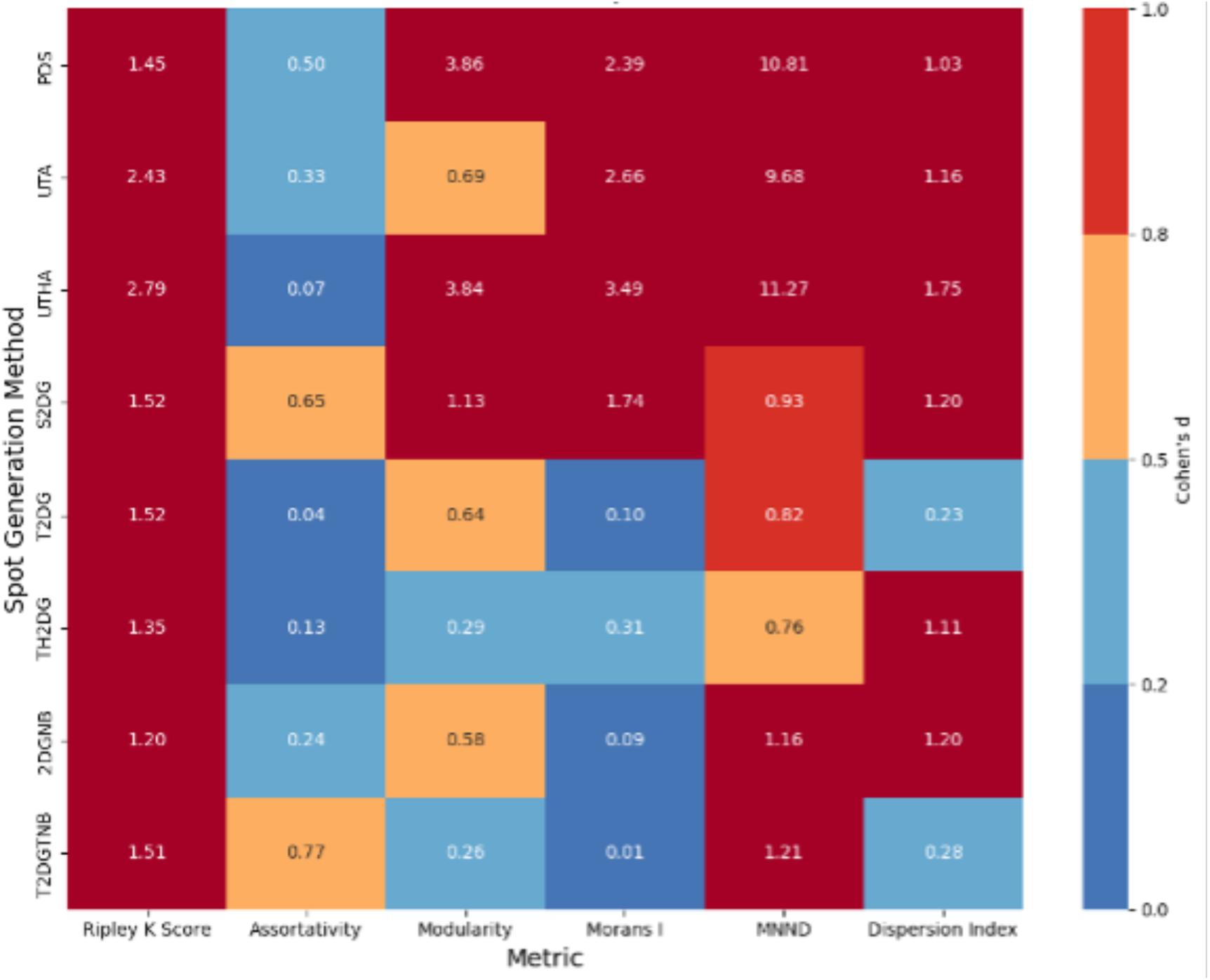
Heatmap displaying Cohen’s D values for clustering metrics across synthetic spatial distribution models. Metrics include Ripley’s K Score, Assortativity, Modularity, Moran’s I, Mean Nearest Neighbor Distance (MNND), and Dispersion Index. Cohen’s D values quantify the effect size between CSR and other spatial distributions: negligible (D<0.2, dark blue), small (0.2<D < 0.5, light blue), medium (0.5 ≤ D < 0.8, orange), and large (D ≥ 0.8, red). Ripley’s K Score consistently demonstrates large effect sizes across most spatial distribution models, indicating substantial differences from CSR. MNND also shows significant differences but is sensitive to dispersion in certain models (e.g., PDS). Other metrics, such as Assortativity and Moran’s I, exhibit moderate to small effect sizes for specific distributions, highlighting their limitations in detecting clustering changes robustly. These results support Ripley’s K Score as a reliable metric for distinguishing centromere clustering patterns.

**Supplementary Figure 2.**
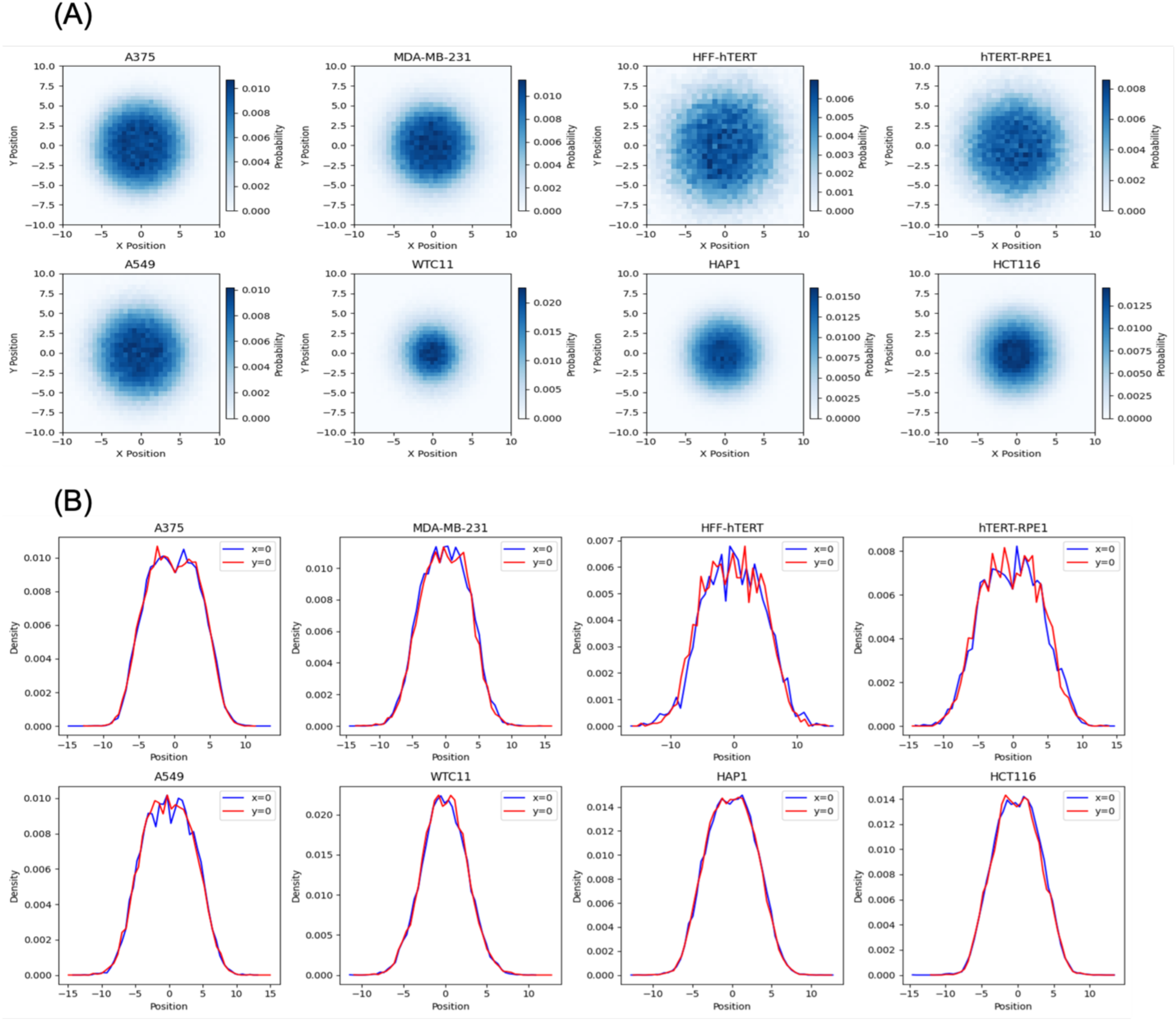
Spatial distribution of standardized centromere locations across eight wild-type cell lines. (A) Overlay of nucleus-centered spot locations from HCT116 (colon), A375 (melanoma), MDA-MB-231 (breast), HFF-hTERT (fibroblast), hTERT-RPE1 (retinal), A549 (lung), HAP1 (myeloid), and WTC11 (embryonic stem cells) (A) 2D histogram showing the distribution of CENP-C spots relative to nuclear center (0,0), revealing a doughnut-shaped pattern in all cell lines. (B) Line plot analysis at x=0 and y=0 demonstrating lower spot density at the nuclear center and higher density between center and nuclear edge.

**Table 1.** Spot generation models abbreviations.

| **Name** | **Abbreviation** |
| --- | --- |
| Complete Spatial Randomness | CSR |
| Poisson-Disk Sampling | PDS |
| Uniformly distributed with Two Adjacent spots | UTA |
| Uniformly distributed with Three Adjacent spots | UTHA |
| Single Two-Dimensional Gaussian | S2DG |
| Two Two-Dimensional Gaussian | T2DG |
| Three Two-Dimensional Gaussian | TH2DG |
| Two-Dimensional Gaussian with a Nuclear Body | 2DGNB |
| Two Two-Dimensional Gaussian with Two Nuclear Bodies | T2DGTNB |
| Real Data | M0 |
| Uniformly Distributed Spots | M1 |
| Cell-Shaped 2D Gaussian | M2 |
| Radial Gaussian Distribution using Ripley's K Function | M3 |
| Bayesian Estimation of Radially Shifted Gaussian using Spot Coordinates | M4 |
| Bayesian Estimation of Radially Shifted Gaussian using Spot Distances | M5 |

**Supplementary Table 1.** The sources of cell lines, culture conditions, media compositions and relevant references of respective culture protocols for the eight cell lines.

| *Metric* | *Reference Group* | *Comparison Group* | *Stat* | *p-value* | *Corrected p-value* | *Significant* |
| --- | --- | --- | --- | --- | --- | --- |
| *Ripley's K Score* | CSR | UTHA | 19621 | 4.37E-303 | **5.8254e-303** | TRUE |
| *Ripley's K Score* | CSR | 2DGNB | 625446 | 0 | **0.0000e+00** | TRUE |
| *Ripley's K Score* | CSR | PDS | 930000 | 5.86E-298 | **6.6992e-298** | TRUE |
| *Ripley's K Score* | CSR | UTA | 33428 | 4.85E-286 | **4.8478e-286** | TRUE |
| *Ripley's K Score* | CSR | T2DGTNB | 893178 | 0 | **0.0000e+00** | TRUE |
| *Ripley's K Score* | CSR | S2DG | 525861.5 | 0 | **0.0000e+00** | TRUE |
| *Ripley's K Score* | CSR | T2DG | 1055834 | 0 | **0.0000e+00** | TRUE |
| *Ripley's K Score* | CSR | TH2DG | 1206212.5 | 0 | **0.0000e+00** | TRUE |
| *Assortativity* | CSR | UTHA | 536942 | 0.00422625 | **4.8300e-03** | TRUE |
| *Assortativity* | CSR | 2DGNB | 4287014 | 9.58E-14 | **1.9162e-13** | TRUE |
| *Assortativity* | CSR | PDS | 622797 | 1.92E-21 | **5.1138e-21** | TRUE |
| *Assortativity* | CSR | UTA | 593698 | 3.99E-13 | **6.3810e-13** | TRUE |
| *Assortativity* | CSR | T2DGTNB | 2756891 | 2.25E-121 | **1.7962e-120** | TRUE |
| *Assortativity* | CSR | S2DG | 3160118 | 2.72E-82 | **1.0865e-81** | TRUE |
| *Assortativity* | CSR | T2DG | 4934849 | 0.49622343 | 4.96E-01 | FALSE |
| *Assortativity* | CSR | TH2DG | 4542913 | 1.81E-06 | **2.4093e-06** | TRUE |
| *Modularity* | CSR | UTHA | 5385 | 0 | **0.0000e+00** | TRUE |
| *Modularity* | CSR | 2DGNB | 3349885 | 1.47E-66 | **2.3572e-66** | TRUE |
| *Modularity* | CSR | PDS | 997501 | 0 | **0.0000e+00** | TRUE |
| *Modularity* | CSR | UTA | 304291 | 6.94E-52 | **9.2591e-52** | TRUE |
| *Modularity* | CSR | T2DGTNB | 4236213 | 1.50E-15 | **1.4978e-15** | TRUE |
| *Modularity* | CSR | S2DG | 8020951 | 1.72E-218 | **4.5798e-218** | TRUE |
| *Modularity* | CSR | T2DG | 6760547 | 1.66E-75 | **3.3104e-75** | TRUE |
| *Modularity* | CSR | TH2DG | 6028129 | 6.75E-27 | **7.7162e-27** | TRUE |
| *Moran's I* | CSR | UTHA | 7523 | 0 | **0.0000e+00** | TRUE |
| *Moran's I* | CSR | 2DGNB | 4882722 | 0.22062501 | 2.21E-01 | FALSE |
| *Moran's I* | CSR | PDS | 972447 | 4.71E-293 | **9.4114e-293** | TRUE |
| *Moran's I* | CSR | UTA | 21241 | 7.05E-301 | **1.8803e-300** | TRUE |
| *Moran's I* | CSR | T2DGTNB | 5407525 | 2.08E-05 | **2.3757e-05** | TRUE |
| *Moran's I* | CSR | S2DG | 9168219 | 0 | **0.0000e+00** | TRUE |
| *Moran's I* | CSR | T2DG | 5543492 | 1.38E-08 | **1.8349e-08** | TRUE |
| *Moran's I* | CSR | TH2DG | 6712341 | 1.57E-71 | **2.5143e-71** | TRUE |
| *MNND* | CSR | UTHA | 1000000 | 0 | **0.0000e+00** | TRUE |
| *MNND* | CSR | 2DGNB | 8778986 | 0 | **0.0000e+00** | TRUE |
| *MNND* | CSR | PDS | 0 | 0 | **0.0000e+00** | TRUE |
| *MNND* | CSR | UTA | 1000000 | 0 | **0.0000e+00** | TRUE |
| *MNND* | CSR | T2DGTNB | 8609522 | 0 | **0.0000e+00** | TRUE |
| *MNND* | CSR | S2DG | 8118659 | 1.03E-232 | **1.3698e-232** | TRUE |
| *MNND* | CSR | T2DG | 7673964 | 1.24E-171 | **1.4190e-171** | TRUE |
| *MNND* | CSR | TH2DG | 7388535 | 2.34E-137 | **2.3425e-137** | TRUE |
| *Dispersion Index* | CSR | UTHA | 93180 | 7.60E-218 | **1.5205e-217** | TRUE |
| *Dispersion Index* | CSR | 2DGNB | 9094784 | 0 | **0.0000e+00** | TRUE |
| *Dispersion Index* | CSR | PDS | 764597 | 2.62E-93 | **3.4939e-93** | TRUE |
| *Dispersion Index* | CSR | UTA | 200042 | 2.33E-119 | **3.7284e-119** | TRUE |
| *Dispersion Index* | CSR | T2DGTNB | 4461777 | 1.90E-08 | **2.1661e-08** | TRUE |
| *Dispersion Index* | CSR | S2DG | 9583300 | 0 | **0.0000e+00** | TRUE |
| *Dispersion Index* | CSR | T2DG | 5230485 | 0.01607436 | **1.6074e-02** | TRUE |
| *Dispersion Index* | CSR | TH2DG | 8142653 | 2.82E-236 | **7.5134e-236** | TRUE |

**Supplementary Table 2.** Statistical testing results for pairwise Mann-Whitney U tests, corrected using the Benjamini-Hochberg False Discovery Rate (BH-FDR) method, comparing clustering metrics calculated for CSR and all other spot generation methods.

| *Metric* | *Distribution* | *Percent Change (%)* |
| --- | --- | --- |
| *Ripley's K Score* | CSR | 59.5567608 |
| *Ripley's K Score* | PDS | inf |
| *Ripley's K Score* | UTA | 12.0172256 |
| *Ripley's K Score* | UTHA | 3.14291737 |
| *Ripley's K Score* | S2DG | 2.83261342 |
| *Ripley's K Score* | T2DG | 1.36536147 |
| *Ripley's K Score* | TH2DG | 1.86411296 |
| *Ripley's K Score* | 2DGNB | 1.59446813 |
| *Ripley's K Score* | T2DGTNB | 4.94364368 |
| *Assortativity* | CSR | 682.231385 |
| *Assortativity* | PDS | 49726.8791 |
| *Assortativity* | UTA | 44496.7239 |
| *Assortativity* | UTHA | 857.562303 |
| *Assortativity* | S2DG | 339.190031 |
| *Assortativity* | T2DG | 683.523035 |
| *Assortativity* | TH2DG | 554.275336 |
| *Assortativity* | 2DGNB | 406.583787 |
| *Assortativity* | T2DGTNB | 280.373907 |
| *Modularity* | CSR | 52.4011886 |
| *Modularity* | PDS | 54.152431 |
| *Modularity* | UTA | 100 |
| *Modularity* | UTHA | 39.8715057 |
| *Modularity* | S2DG | 55.0145623 |
| *Modularity* | T2DG | 50.1452432 |
| *Modularity* | TH2DG | 52.453178 |
| *Modularity* | 2DGNB | 50.1102691 |
| *Modularity* | T2DGTNB | 51.4916329 |
| *Moran's I* | CSR | 17.7412248 |
| *Moran's I* | PDS | 17.6846799 |
| *Moran's I* | UTA | 132.437197 |
| *Moran's I* | UTHA | 13.5630049 |
| *Moran's I* | S2DG | 15.6657324 |
| *Moran's I* | T2DG | 14.8906965 |
| *Moran's I* | TH2DG | 15.3107854 |
| *Moran's I* | 2DGNB | 14.7192826 |
| *Moran's I* | T2DGTNB | 15.4520555 |
| *MNND* | CSR | 77.2792286 |
| *MNND* | PDS | 32.7068625 |
| *MNND* | UTA | 1160.51333 |
| *MNND* | UTHA | 273.062842 |
| *MNND* | S2DG | 69.7615691 |
| *MNND* | T2DG | 72.571224 |
| *MNND* | TH2DG | 72.5743327 |
| *MNND* | 2DGNB | 75.9056292 |
| *MNND* | T2DGTNB | 84.1758623 |
| *Dispersion Index* | CSR | 2.34686669 |
| *Dispersion Index* | PDS | 2.25438507 |
| *Dispersion Index* | UTA | 38.722864 |
| *Dispersion Index* | UTHA | 2.30607869 |
| *Dispersion Index* | S2DG | 3.97306092 |
| *Dispersion Index* | T2DG | 2.79085127 |
| *Dispersion Index* | TH2DG | 2.862863 |
| *Dispersion Index* | 2DGNB | 2.36469957 |
| *Dispersion Index* | T2DGTNB | 3.02788452 |

**Supplementary Table 3.**
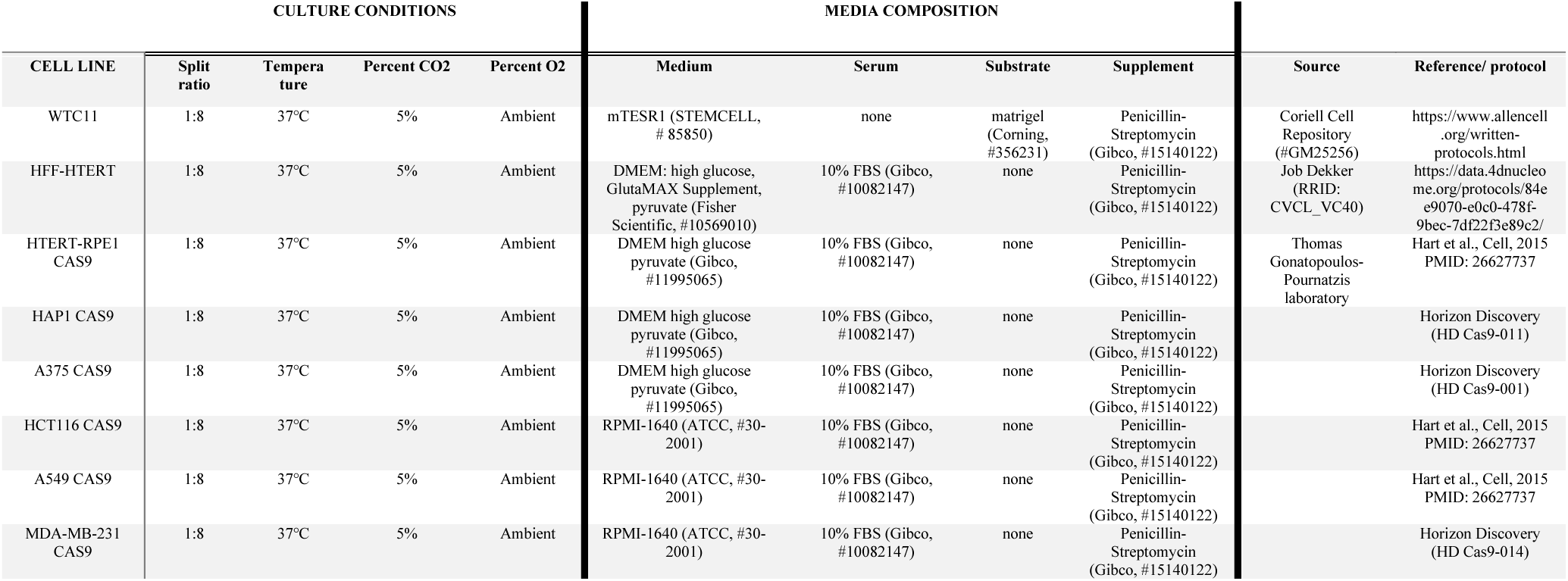
The percent change calculated as average value for each metric for up to 30 spots removed from the initial 46 spots compared to the value with 46 spots.

